# Deep learning recognises antibiotic modes of action from brightfield images

**DOI:** 10.1101/2025.03.30.645928

**Authors:** Daniel Krentzel, Kelvin Kho, Julienne Petit, Nassim Mahtal, Jeanne Chiaravalli, Spencer L. Shorte, Anne Marie Wehenkel, Ivo G. Boneca, Christophe Zimmer

## Abstract

Antimicrobial resistance is a growing public health threat predicted to cause up to 10 million deaths a year by 2050. To circumvent existing bacterial resistance mechanisms, discovering antibiotics with novel modes of action (MoAs) is crucial. While growth inhibition assays can robustly identify antibiotic molecules, they miss promising compounds with subinhibitory phenotypes and do not inform on drug MoA. Microscopy-based cytological profiling of drug-treated bacteria with hand-crafted image descriptors or more recently deep learning (DL) provides complementary information on the MoA. However, current approaches are limited by the need for fluorescent labelling and drug exposure at inhibitory concentrations. It also remains unclear if cytological profiling enables the detection of drugs with novel MoAs. Here, we demonstrate an approach based on supervised DL to identify antibiotic drug MoA from microscopy images without fluorescent labelling. We train a convolutional neural network to predict treatment conditions from brightfield images of *Escherichia coli* exposed to reference drugs covering multiple MoAs. Our method can detect drug exposure at subinhibitory concentrations and distinguishes individual drug treatments with high accuracy (86.2%). The learned representations implicitly capture MoA-specific phenotypes, enabling perfect MoA recognition (100%) without retraining using a model trained on only 644 images. Our approach can identify the MoA of previously unseen drugs with good accuracy (77.8±3.3%), as long as the MoA is represented by at least one of the training compounds. Finally, we show that our approach can detect if a drug has a novel MoA with an area under the curve above 0.75 for five out of six MoAs, facilitating microscopy-based identification of novel classes of antibiotics. Our methods and results pave the way towards an automated pipeline for antibiotic drug discovery based on imaging and DL.

## Introduction

Antimicrobial resistance (AMR) caused 1.27 million deaths in 2019 (1) with death tolls estimated to rise to as much as 10 million deaths a year by 2050 (2). During the last 40 years, very few antibiotics with novel modes of action (MoAs) have been discovered. New resistance mechanisms of various pathogenic bacteria have thus remained unchecked, thereby increasingly rendering the existing arsenal of antibiotics ineffective (3).

Phenotypic drug discovery (PDD) is an attractive approach to discover antibiotics with new MoAs, since no prior target hypothesis is required. A typical read-out in PDD screens is optical density (OD) of compound-treated bacterial broths, which serves as a proxy for bacterial cell density. While this read-out allows to identify compounds that inhibit bacterial growth at concentrations exceeding the minimum inhibitory concentration (MIC), it precludes the identification of antibacterial compounds at sub-MIC concentrations that could be optimised by medicinal chemistry or be effective in combination therapies. Moreover, these single read-out screens cannot provide information on how an antibacterial compound inhibited bacterial growth, i.e. the MoA (4).

To overcome these limitations, recent antibacterial screening campaigns (5) have drawn inspiration from eukaryotic PDD approaches using microscopy images as read-outs (6). Nonejuie *et al.* (7) and Zoffmann *et al.* (8) acquired fluorescence microscopy images of bacteria and extracted hand-crafted quantitative descriptors (features) from segmented bacterial cells to perform bacterial cytological profiling (BCP). Importantly, compounds with the same MoA induced similar phenotypic changes, thus enabling the computational clustering of compounds by MoA directly from microscopy images. However, a limitation of early BCP is its reliance on cell segmentation, an error-prone step especially at high cell densities, and the potentially insufficient or unspecific information contained in hand-crafted features (4). Deep learning (DL) approaches, in contrast, hold the potential to extract highly specific phenotypic features from images in an end-to-end manner without segmentation (4). Prior work on microscopy images of eukaryotic cells labelled with multiple fluorescent dyes (Cell Painting) (9) has shown that features learned by a convolutional neural network (CNN) can outperform handcrafted features in discriminating and associating different and similar treatment conditions respectively (10). Recently, a CNN trained on over 5,000 images has been shown to successfully classify the MoA of antibiotics at concentrations of 1-5×MIC from multi-channel images of fluorescently stained *Mycobacterium tuberculosis* bacteria (11).

Despite these encouraging efforts, several limitations and questions remain to be addressed before DL analysis of high-throughput microscopy images can be leveraged efficiently in future screens to discover antibiotic compounds with novel MoAs. First, it is unclear if antibiotic compounds can be identified from images at sub-MIC concentrations, which would greatly improve the sensitivity of large-scale screens that typically test thousands of compounds at a single concentration. Second, it remains to be determined to what extent unseen compounds can be assigned to their correct MoA and, critically, whether compounds with novel MoAs can be detected. Third, fluorescent labelling is expensive, significantly adds complexity to experimental protocols, can introduce batch-to-batch variability, and may interfere with the molecular mechanism of compounds. Whether fluorescent labelling can be dispensed with in favour of brightfield (BF) images in phenotypic screens of bacteria, as recently shown in eukaryotic cells (12), is unclear, since bacterial cells are roughly an order of magnitude smaller, potentially limiting the range of visible morphological phenotypes. Finally, the performance of DL typically comes at a steep cost in training data, as e.g. ∼200 fields of view (FOV) per compound were used to train a CNN in (11), imposing limits to scaling up this approach to thousands of compounds and calling for more efficient approaches.

Here, using *Escherichia coli*, we develop and validate a DL approach to extract bacterial phenotypic features end-to-end directly from BF images, thus dispensing with the need for fluorescent staining. We show that from these BF images alone our model can reveal alterations induced by antibiotic compounds at concentrations well below the MIC and accurately recognise the MoA of compounds when trained on as few as seven images per well. Moreover, our approach can recognise the MoA of unseen compounds by comparing to reference antibiotics of the same MoA, and determine if a compound has a novel MoA, again from BF images only. Our results lay the foundations for a time- and cost-effective pipeline to identify chemical compounds with potentially novel antibiotic MoAs.

## Results

### Collecting a high-throughput image dataset of bacterial phenotypes

We collected a high-throughput imaging dataset of *E. coli* bacteria exposed to 22 widely used reference drugs. We labelled each of these drugs using only one of eight classes of MoA based on information from the literature (**Table 1**). Our reference drugs included seven inhibitors of cell wall synthesis targeting penicillin binding proteins (PBPs), which we labelled based on their primary target as inhibitors of PBP1A/B, PBP2 or PBP3 (**Supp.Table 1**); four β-lactamase inhibitors shown to have moderate to weak activity towards these PBPs; four antibiotics targeting ribosomes; three targeting DNA gyrase; two targeting membrane integrity; one targeting RNA polymerase, and one targeting DNA synthesis. Relebactam is the only drug for which no targets had been reported in *E. coli*. As it belongs to the family of diazabicyclooctanes (DBOs) that includes avibactam - a PBP2 inhibitor, we labelled relebactam as a PBP2 inhibitor. For each of these drugs, we determined the half maximal inhibitory concentration (IC_50_) (**Methods**), which ranged from 5.7 ng/ml to 64 μg/ml (**Table 1**). We distributed these antibiotics onto four 96-well plates with randomised layouts (**Supp. Fig. 1**) using an acoustic liquid handler (**Methods**) to expose bacteria to one of four subinhibitory concentrations (0.125×IC_50_, 0.25×IC_50_, 0.5×IC_50_, IC_50_). Then, we grew bacteria to exponential phase and seeded them onto the four 96-well plates containing the previously distributed antibiotics (**Methods**). After a 2h15min incubation followed by bacterial membrane staining with FM4-64FX for 15 min, we fixed bacteria with a combination of paraformaldehyde and glutaraldehyde whereupon we stained DNA with Hoechst 33342. Next, we acquired high-throughput images (**Methods**) in three channels (BF, FM4-64 and Hoechst) using a high-content screening system in non-confocal mode with a 63× water-immersion objective (**Fig. 1a, 2a**).

**Fig. 1.**
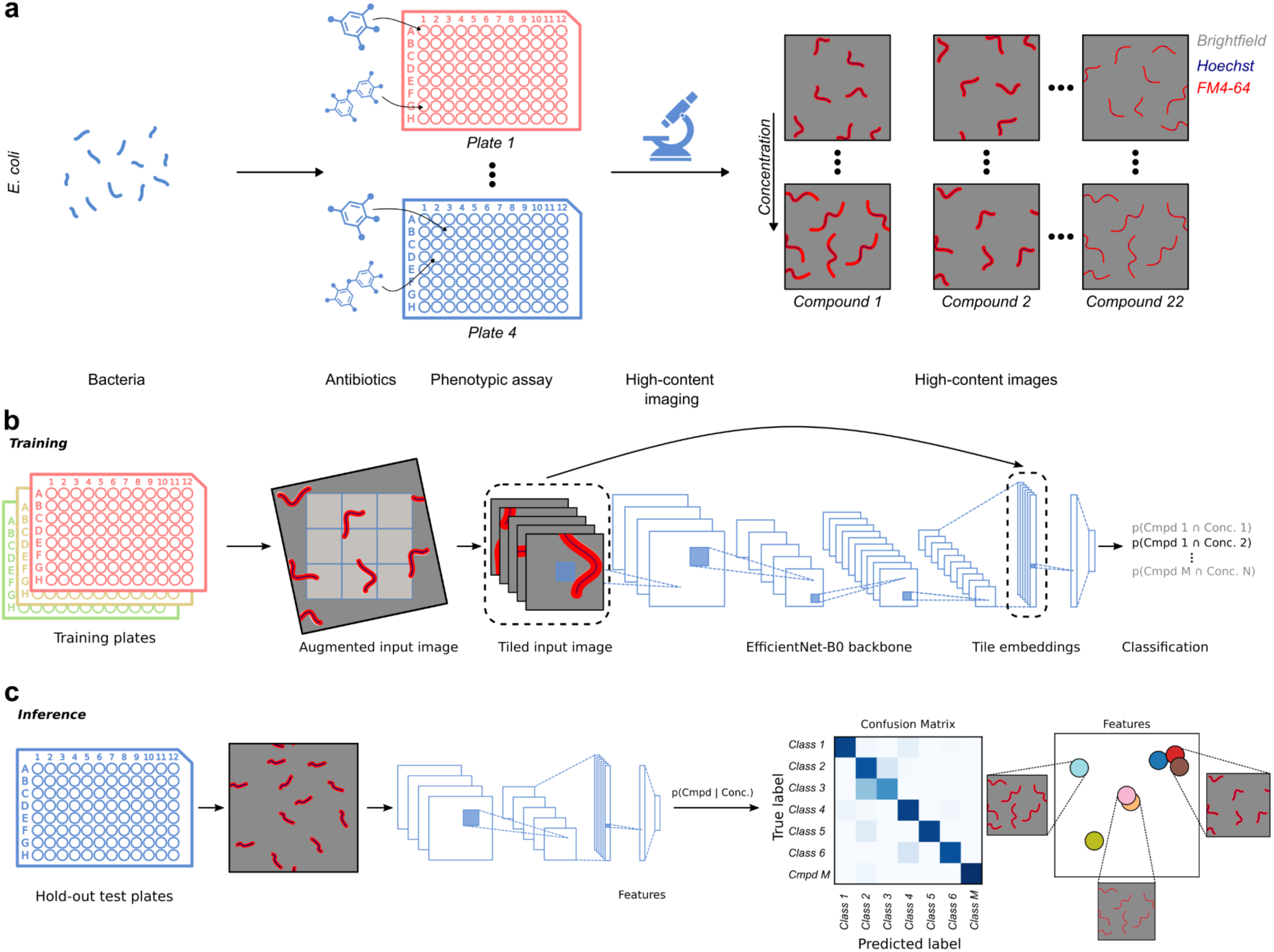
Data acquisition, model training and inference. (**a**) *E. coli* bacteria were grown to exponential phase in culture, antibiotics were distributed onto four 96-well plates at four different concentrations with an acoustic liquid handler and bacteria dispatched onto the well plates. Bacteria were then grown with antibiotics for 2h15min and fixed. Plates were imaged on a high-content microscope with a 63× water-immersion objective in non-confocal mode with three channels (BF, FM4-64FX and Hoechst). (**b**) The model was trained on high-throughput images from three out of four plates to predict the combination of antibiotic and concentration with a cross-entropy loss. Images were first passed through a data augmentation pipeline and then tiled. Each tile was then passed through the same EfficientNet-B0 backbone whereupon embedding vectors from each image tile were aggregated to obtain a global embedding vector. (**c**) Testing was performed on a hold-out test plate with predictions computed at a given concentration and embedding vectors visualised in 2D with UMAP.

**Table 1:** Antibiotics used in this study, their MoA according to literature and their potency against *E. coli*.

| MoA |  | Antibiotic | IC<sub>50</sub> (µg/ml) | MoA reference |
| --- | --- | --- | --- | --- |
| Cell wall | PBP1A/B | Cefsulodin | 23.77 | (23,32) |
|  |  | Penicillin G | 25.65 | (23,33) |
|  |  | Sulbactam | 15.62 | (24) |
|  | PBP2 | Avibactam | 19.20 | (33) |
|  |  | Mecillinam | 0.3009 | (23,32,34) |
|  |  | Meropenem | 0.1085 | (33,35) |
|  |  | Clavulanate | 17.24 | (24) |
|  |  | Relebactam | 64 | N/A for *E. coli* |
|  | PBP3 | Aztreonam | 0.01568 | (33) |
|  |  | Cefepime | 0.007517 | (33,36) |
|  |  | Ceftriaxone | 0.004 | (33) |
| Ribosome |  | Chloramphenicol | 3.079 | (37) |
|  |  | Clarithromycin | 22.24 | (38) |
|  |  | Doxycycline | 0.3703 | (39) |
|  |  | Kanamycin | 2.529 | (40) |
| DNA gyrase |  | Ciprofloxacin | 0.005723 | (41) |
|  |  | Levofloxacin | 0.01089 | (42) |
|  |  | Norfloxacin | 0.02538 | (43) |
| RNA polymerase |  | Rifampicin | 3.358 | (44) |
| DNA synthesis |  | Trimethoprim | 0.1335 | (45) |
| Membrane integrity |  | Colistin | 0.6465 | (46) |
|  |  | Polymyxin B | 1.166 | (47) |

### Deep learning model and training strategy

We then designed a DL model (**Supp. File 1**) to map images of antibiotic-treated bacteria to chemical perturbations (**Methods**). We based our model on the CNN architecture EfficientNet-B0 (13), which was shown to achieve state-of-the-art performance on established image classification benchmarks with much fewer parameters than more widely used architectures like ResNet-50 (14) (e.g. 5.3 million parameters vs. 26 million parameters and 76.3% vs. 76.0% accuracy on ImageNet classification (13,15)), thereby keeping memory requirements and computation time moderate.

As input data we used images of entire FoVs, of 2,160×2,160 pixels (corresponding to an area of 139.32×139.32 μm). Images were passed through a data augmentation pipeline and divided into nine tiles which were processed in parallel using the same convolutional filters. Tile-wise embedding vectors were aggregated before the final classification layer (**Fig. 1b**; **Methods**), thus diminishing the influence of tiles without bacteria on model training.

We trained our model to perform image classification, where each distinct antibiotic drug treatment, i.e. each combination of antibiotic and concentration (e.g. “aztreonam” and “0.25×IC_50_”) was considered as a separate class, resulting in a total of 89 classes (4 concentrations × 22 compounds + 1 control) (**Methods**). In order to reduce our model’s sensitivity to possible technical confounders (e.g. plate-specific image features and well position-dependent effects independent of the drug-induced phenotypes) and increase its robustness to biological variability, we used three biological replicates where bacteria were cultured independently and grown on different days with randomised well positions (**Supp.** Fig. 1) to train a single model. Critically, to avoid biases and spurious correlations between training and testing data that could lead to an overestimation of classification performance, we evaluated each model’s classification performance on a biological replicate, using a fourth hold-out test plate with a different randomised layout.

### Deep learning accurately recognises antibiotic mode of action from images

We first asked if and to what extent our model can predict the combination of both drug and concentration from images alone, considering all the available imaging modalities (BF, Hoechst, FM4-64FX). This is a very difficult task, since our treatments include multiple compounds with the same targets (**Table 1**) and changes in drug concentration are not necessarily expected to induce different phenotypes (e.g. if concentrations are much below or above the IC_50_). Indeed, although some drug treatments led to easily recognisable phenotypes, distinguishing between all treatment conditions only by visual inspection from images remained extremely challenging (**Supp.** Fig. 2). If the model were unable to learn any features specific to the combination of drug and concentration, the expected random classification accuracy would be 1.12% (1/89). We assessed this accuracy on four rotations of the training and testing plates, with confusion matrices shown in **Supp.** Fig. 3. Remarkably, the classification accuracy ranged from 31.3% to 47.4% at the image level, i.e. 28 to 42 times better than random. This indicates that the model extracted information specific to drug-concentration combinations from images, although far from allowing perfect classification.

As the effect of antibiotics is likely to be more pronounced at higher concentrations, we next assessed our model (trained on all drug-concentration combinations) only on test data at the highest drug concentrations, i.e. the IC_50_, and assessed accuracy based only on the predicted antibiotic (thus ignoring the predicted concentration). On these data, our model achieved a very high per-image accuracy of 76.2% to 90.2%, compared to 4.3% (1/23) for random classification (**Supp.** Fig. 4). Per-well accuracies (obtained using the majority of predictions from all images in the same well) were slightly higher, namely 78.3% to 91.3%, indicating that at the IC_50_, the vast majority of the 22 reference drugs used here led to highly specific phenotypes, which our model was able to automatically learn from the training data (**Fig. 2b**).

**Fig. 2.**
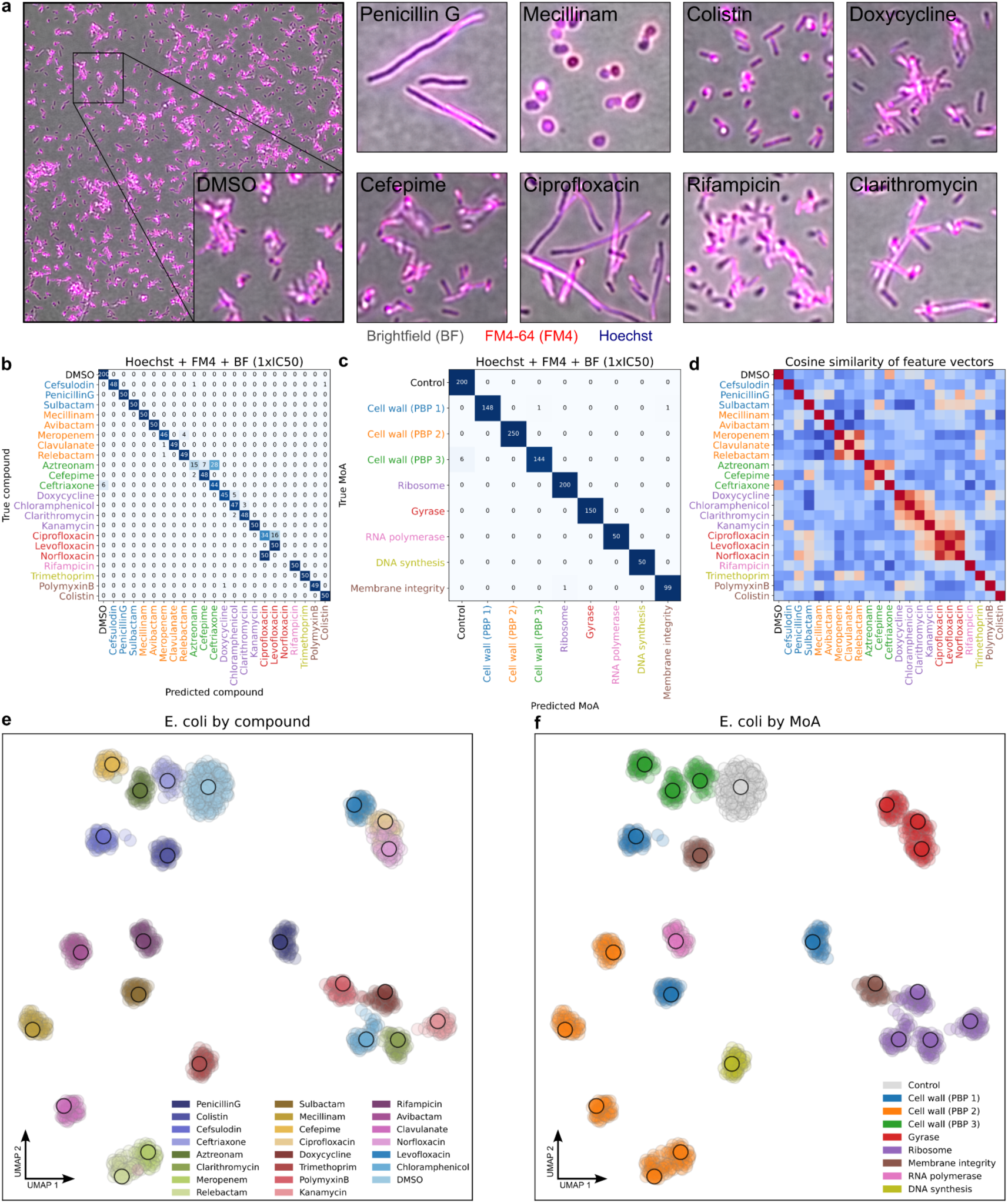
Deep learning recognises compounds and their MoA from *E. coli* images. **(a)** Representative images from the test plate used to generate results with BF shown in grey, FM4-64FX in red and Hoechst in blue. See **Supp.** Fig. 2a-f for more examples. (**b**) Confusion matrix showing predictions from high-throughput images of bacteria treated with the same relative concentration (IC50) from a model trained on all imaging channels and concentrations. (**c**) Confusion matrix by MoA was obtained by pooling predictions of compounds belonging to the same MoA (**Table 1**). (**d**) Cosine similarities between embedding vectors obtained from bacterial high-throughput images were computed on median feature vectors. High cosine similarity (red) indicates phenotypic similarity between conditions and tends to occur for antibiotics belonging to the same MoA (e.g. ciprofloxacin, norfloxacin and moxifloxacin target the DNA Gyrase). (**e**) UMAP was used to project high-dimensional feature vectors to 2D. Small dots correspond to individual high-throughput images and large dots to median feature vectors. Dots are coloured by antibiotic compounds. (**f**) UMAP with dots coloured by MoA shows that antibiotics belonging to the same MoA tend to co-localise in latent space (e.g. PBP3 inhibitors in green and DNA gyrase inhibitors in red).

Nevertheless, prediction of antibiotic identity was not perfect, as the model still tended to confuse certain compounds. Inspection of individual hold-out test replicates revealed that the model tended to systematically confuse a subset of drugs (**Fig. 2b**, **Supp.** Fig. 4). For example, aztreonam was confused with cefepime, cefsulodin or ceftriaxone for 3 out of 4, and relebactam was confused with meropenem in 2 out of 4 pairs of model and test data, and norfloxacin was systematically mistaken for ciprofloxacin in one hold-out test dataset. We noted that the confused compounds tended to have similar MoAs (**Table 1**). Moreover, when computing cosine distances between median embeddings for all pairs of antibiotics, we could observe clustering by MoA (**Fig. 2d**), as also illustrated by UMAPs (**Fig. 2e,f**), suggesting that our model extracted similar feature vectors from images for compounds targeting the same MoA despite only having been trained on drug labels.

This observation encouraged us to assess our model’s ability to recognise the MoA, as opposed to the identity of the drugs themselves. For this purpose, we assigned each of the reference drugs to one out of six MoAs or target classes, namely: cell wall, ribosome, DNA gyrase, RNA polymerase, DNA synthesis and membrane integrity (**Table 1**). Because half of the antibiotics were cell wall inhibitors (11 out of 22), we further subdivided this class by their main target proteins, namely PBP1A/B, 2 and 3 (**Table 1**), thereby leading to eight classes of targets with one to five antibiotics per target class. We then assessed the model’s performance at recognising the MoA of each drug by pooling drugs and predictions according to their known MoA. Note that this did not involve any retraining on MoAs as we still used the model trained on combinations of antibiotics and concentrations. Remarkably, our model achieved an accuracy of 84.1% to 99.3% in classifying drug MoA at the image-level and of 100% for three out of four replicates at the well-level (on the remaining plate the classification accuracy was 88.9%), indicating almost perfect recognition of MoA from images (**Fig. 2c**).

Thus, our data shows that images of bacteria treated with reference antibiotics at IC_50_ concentrations contain highly specific information about drug MoA, allowing accurate MoA recognition from images alone. Moreover, our results indicate that established CNN architectures are well suited to extract this information in an end-to-end fashion when trained on drug labels alone, without the need for handcrafted features, cell segmentation and *a priori* knowledge of drug MoAs.

### Brightfield images alone allow accurate recognition of drug MoA

In contrast to BF, fluorescent labelling of various bacterial cell compartments requires selection of dyes with compatible excitation and emission spectra, as well as the development of appropriate staining protocols. We therefore asked if all three imaging channels, namely BF, FM4-64FX and Hoechst (**Fig. 3a**) were necessary to accurately distinguish between antibiotic drugs or MoA. To quantify this, we trained several CNN models as described previously, but with all possible combinations of the three imaging channels as input and assessed classification accuracy on the hold-out test data using the same channels. In some cases, the accuracy varied significantly between the four hold-out plates, with accuracies ranging from 65.2% to 95.7% for the models trained only on the FM4-64FX channel. We speculate that these differences could be due to variations in staining efficiency between plates. The median accuracy over the four replicates was highest for the models trained and tested only on the two fluorescent channels and all three channels together (**Fig. 3b, Supp.** Fig. 5). To our surprise, however, ignoring one or even two of the channels in training and testing resulted in only small reductions of median accuracy when compared to the differences between plates (**Supp.** Fig. 6). This suggests that the three channels contained largely redundant information. Strikingly, when considering the BF channel alone (**Fig. 3d**), the median accuracy was 82.6%, only 4.4% below the three-channel accuracy, and was more consistent across plates (accuracies ranged from 73.9% to 82.6% vs. 78.3% to 91.3%). Pooling the predictions from the BF- only model by MoA led to perfect MoA classification (**Fig. 3c,e**) and feature vectors obtained from the BF-only model also clustered by MoA (**Fig. 3f**).

**Fig. 3.**
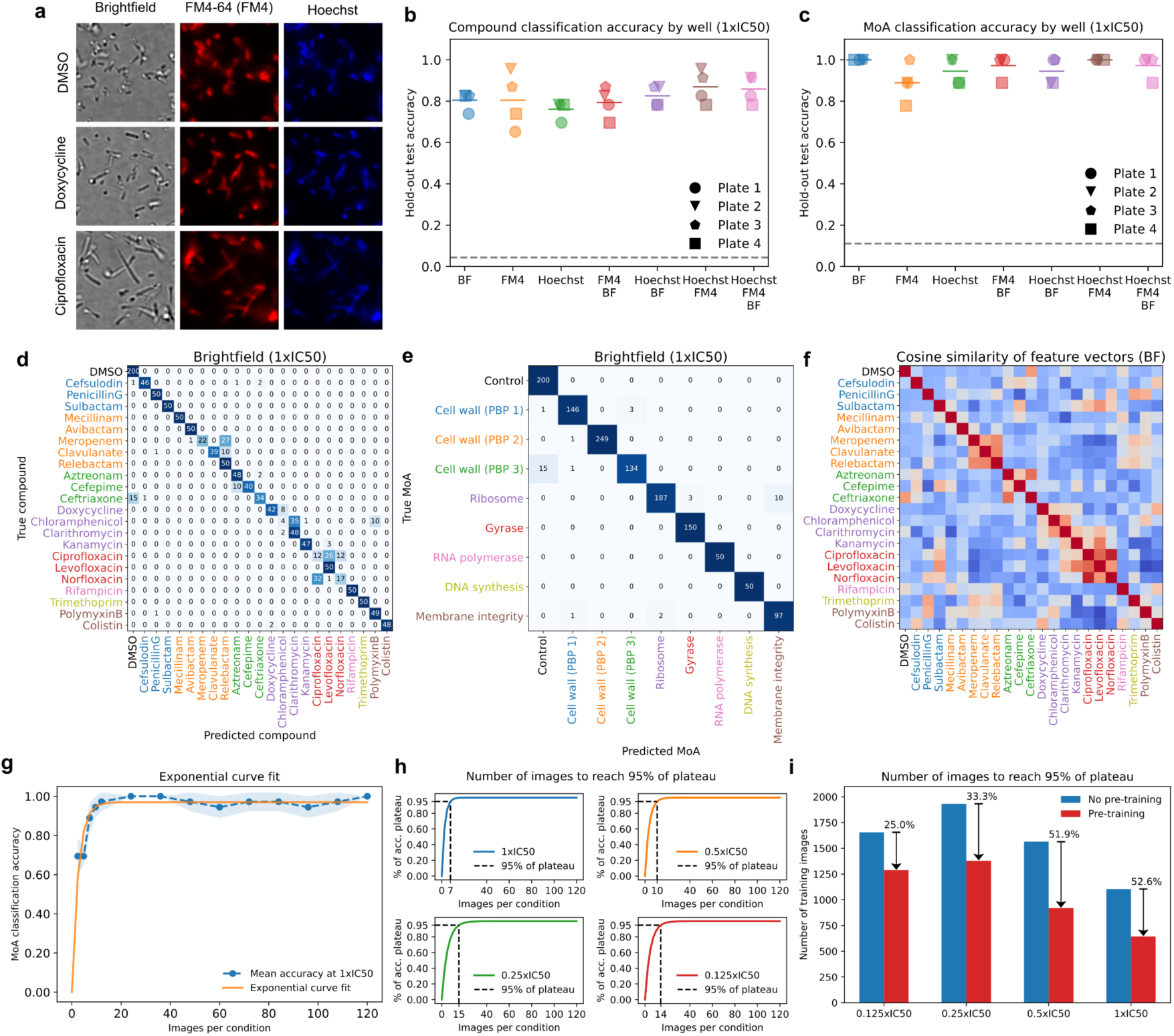
Classification performance for different numbers of imaging channels and training images. **(a)** Example images split by imaging channel. (**b**) Models were trained in a cross-validation experiment with three plates using different combinations of imaging channels as input and tested on a hold-out test plate. Per-well compound classification accuracies were obtained by aggregating predictions from all images acquired in the same well of bacteria treated at the same relative concentration (IC50). The dashed line indicates random classification. (**c**) Per-well MoA classification accuracies were obtained by pooling per-well compound predictions. Models trained on BF images alone perform on par with models trained on Hoechst and BF images. The dashed line indicates random classification. (**d**) Confusion matrix showing predictions from high-throughput images of bacteria treated with the same relative concentration (IC50) from a model trained only on BF images. (**e**) Confusion matrix by MoA was obtained by pooling predictions of compounds belonging to the same MoA (**Table 1**). (**f**) Cosine similarities between embedding vectors obtained from only BF images were computed on median feature vectors. High cosine similarity (red) indicates phenotypic similarity between conditions and is observed for antibiotics belonging to the same MoA (e.g. meropenem, clavulanate and relebactam target PBP2). (**g, h**) MoA classification accuracy as function of the number of training images. The number of images required to reach 95% MoA classification accuracy was estimated by fitting a two-parameter exponential function to averaged MoA classification curves with an example shown in (**g**) for predictions obtained from images of bacteria treated at IC50. (**h**) The number of images per condition required to reach 95% of the accuracy plateau was computed for each concentration showing that between seven and 15 images are required. (**i**) Bars show the number of training images required to reach 95% of the accuracy plateau with (red) or without (blue) pretraining on ImageNet evaluated for different concentrations by cross-validation on four hold-out test plates. The number of required training images ranged from 644 (IC50) to 1,380 (0.25×IC50) with pretraining. All models were trained on BF images only.

These results show that accurate MoA recognition is possible from BF images alone, thereby removing the need for fluorescent labelling. This could drastically simplify experimental protocols and significantly reduce costs in large phenotypic screens.

### CNN learns MoA-specific features from only seven images per condition

A major drawback of using DL algorithms compared to classical analysis methods based on hand-crafted features is the need for large amounts of training data. Assessing the minimum number of images required to train a DL model without losing classification performance can help to optimise and potentially reduce the burden of data collection. We therefore estimated the minimum number of images required to train our CNN without adversely affecting MoA classification and also assessed the impact of pretraining our model on a large public dataset of natural images (ImageNet (15)).

To ascertain the minimum number of required images to robustly distinguish between MoA classes, we trained our CNN using an EfficientNet-B0 convolutional backbone pretrained on ImageNet (**Fig. 1b**) with varying numbers of images per condition ranging from three to 120 and computed mean classification accuracies across four replicates. We observed that classification accuracy stopped increasing with the number of training images and fitted a two-parameter exponential function to determine the accuracy plateau and the number of images required to reach it (within 5%), at four different concentrations (**Fig. 3gh, Methods**). We found that between seven (IC_50_) and 15 images (0.25×IC_50_) per condition (644 and 1,380 images in total) were sufficient to achieve the maximum accuracy. Repeating the analysis with randomly initialised weights (**Fig. 3i**), we found that pretraining on ImageNet data reduced the number of required images by between 25.0% (0.125×IC_50_) and 52.6% (IC_50_).

These results show that near-perfect MoA classification performance can be achieved with only seven images per condition for *E. coli* bacteria exposed to compounds at the IC_50_ concentration and highlight the benefit of using pretrained models to reduce the required training data.

### Detecting drug phenotypes at subinhibitory concentrations

The above results show that DL can reliably extract MoA-specific features from images of bacteria exposed to drugs at IC_50_ concentrations, which for the reference drugs considered here ranged from 5.7 ng/ml to 64 μg/ml (**Table 1**). Visual inspection of drug-treated bacteria shows that the cellular phenotypes tend to be less pronounced at lower concentrations, as expected (**Fig. 4a**). When screening large libraries of compounds, systematic titration to determine the IC_50_ for each compound is often not practical. Instead, chemical screens typically rely on a single concentration, which for most active compounds is generally lower or higher than the inhibitory concentration. We therefore asked if our DL method can detect or even classify drug-induced phenotypes at subinhibitory concentrations, and in particular concentrations at or below the IC_50_. Since at sub-MIC concentrations, new chemical compounds may induce only subtle phenotypic effects before optimisation by medicinal chemistry, the ability to detect such weak phenotypes could increase the sensitivity and hit rate of phenotypic screening for antibiotic drug discovery.

**Fig. 4.**
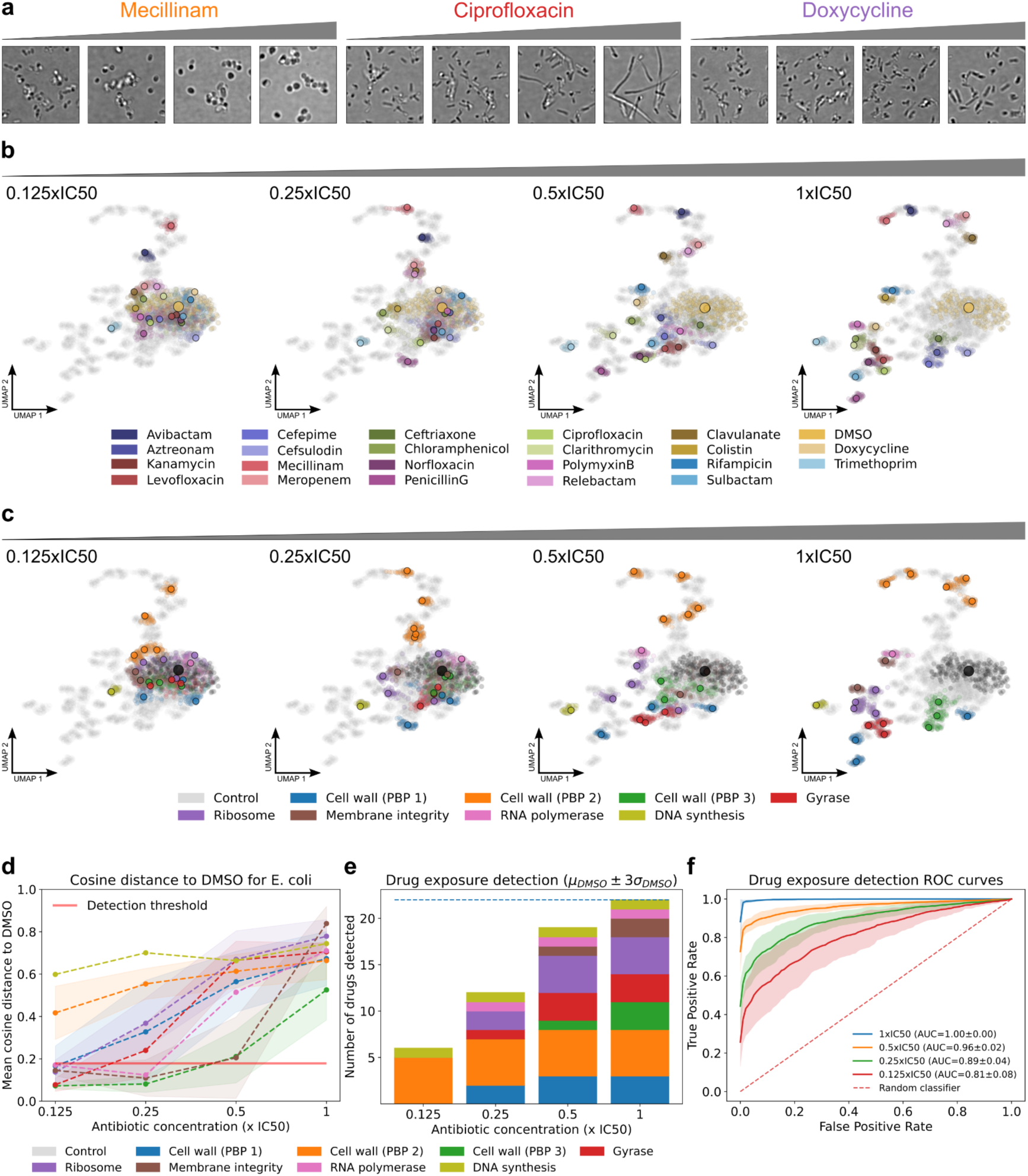
Detecting drug exposure at subinhibitory concentrations. **(a)** Example images of bacteria treated at increasing concentrations show dose-dependent phenotypic changes. (**b**) Feature vectors of bacteria treated at four different concentrations were jointly projected with UMAP and dots corresponding to a given concentration highlighted in each plot. Small dots correspond to individual high-throughput images and larger dots correspond to median feature vectors. Dots are coloured by antibiotics. (**c**) UMAP plots coloured by MoA show that, except for antibiotics targeting PBP2 and DNA synthesis, feature vectors tend to be further away from DMSO as concentration increases. (**d**) The cosine distance between median feature vectors and DMSO was computed for each antibiotic class at different concentrations. Average distances by MoA are shown with dashed lines and shaded areas indicate the standard deviation. A detection threshold (red line) was computed from the intra-class cosine distances of DMSO feature vectors (mean plus three standard deviations). (**e**) Drug exposure detection was determined by evaluating cosine distances between median feature vectors and DMSO with respect to the detection threshold. The number of detected drugs at each relative concentration is shown and bars are coloured by the MoA of the respective antibiotics. The dashed line indicates the overall number of antibiotics. (**f**) ROC curves for drug exposure detection at different concentrations cross-validated on four hold-out test replicates with corresponding AUC values are shown.

We obtained confusion matrices for sub-IC_50_ concentrations and observed that, unsurprisingly, our model’s ability to correctly classify bacterial images by compound increased with higher concentrations (**Supp.** Fig. 7). We then analysed the embeddings computed by our BF-only model (trained on reference antibiotics at multiple concentrations as detailed above) on test data at increasing concentrations, from 0.125×IC_50_ to 1×IC_50_ (**Fig. 4b,c**). For most antibiotics, embeddings were similar to untreated controls (DMSO) at the lowest tested concentration (0.125×IC_50_) and tended to move away from the control and from one another as concentration increased. The exceptions were the PBP2 inhibitors avibactam and mecillinam, as well as the DNA synthesis inhibitor trimethoprim, for which embeddings already localised away from the controls at the lowest concentration, suggesting that these compounds induce distinctive phenotypic changes at concentrations well below inhibitory values. To quantify this observation, we computed the cosine distances between the median embeddings of drug-treated cell images and the DMSO images, at each concentration (**Fig. 4d**). We also defined a threshold distance as the mean plus three standard deviations of the cosine distance between individual DMSO images. Based on this threshold, we found that even at 0.125×IC_50_, compounds from three groups of antibiotics (inhibitors of DNA synthesis, ribosome, PBP1A/B and PBP2) exhibited sufficiently different phenotypes to be detected (**Fig. 4e**). At 0.5×IC_50_, all but two groups of antibiotics (RNA polymerase and PBP3 inhibitors) were distinguished from the negative control.

To assess drug exposure detection based on cosine distance to DMSO at different subinhibitory concentrations, we obtained receiver operating characteristic (ROC) curves cross-validated on four hold-out plates and computed the corresponding area under the curve (AUC) (**Fig. 4f**). We observed that, in line with **Fig. 4e**, drug exposure detection improved with increasing concentrations and that even at the lowest concentration (0.125×IC_50_) drug exposure was robustly detected with an AUC of 0.81±0.08, while at IC_50_ detection was perfect (AUC=1.00).

Overall, our data show that image embeddings learned by our CNN can be used to robustly detect subtle phenotypic changes at subinhibitory concentrations from BF images alone, thereby enabling highly sensitive detection of potentially active compounds in phenotypic screens.

### Recognising known modes of actions of unseen drugs

Our analyses above showed that our model can recognise the identity of a compound or the MoA of several antibiotics after training on independent images obtained with the same antibiotics. However, in the context of a screening campaign, the goal is to determine the MoA of a previously unseen novel drug. Therefore, we next asked if our model can correctly recognise the MoA of an unseen compound, i.e. a compound that was entirely absent from the training data. To address this question, we performed a leave-one-compound-out (LOCO) analysis for compounds sharing a MoA with at least two other compounds.

We retrained our CNN only using BF images by systematically excluding all images of one compound from the training data (**Fig. 5a**). Then, we used this new model to obtain embeddings from images of bacteria exposed to the left-out compound at IC_50_ on a hold-out test plate. Next, we obtained reference embeddings using the same hold-out test plate from images of bacteria exposed to all other compounds at IC_50_. For each FoV, we identified the closest reference compound using a k-NN classifier (k=150 and cosine distance) on embedding vectors (**Fig. 5b**, **Methods**). For each left-out compound, we then assigned the MoA of the reference compound that was the closest for most FoVs. We found that at a concentration of IC_50_, the classification accuracy across plates ranged from 72.2% to 83.3% (**Fig. 5c**). Moreover, we observed that the k-NN classification on embeddings performed better on average than directly using the MoA predictions of our model (75.0±3.5% vs. 63.8±5.4%, **Fig. 5c**). We then evaluated MoA assignment accuracy for a range of concentrations and found that accuracy tended to improve with higher concentration (**Fig. 5d**). Interestingly, drugs targeting PBP2 were perfectly classified across the entire range of concentrations, indicating that these compounds induce highly distinctive phenotypes even at low concentrations.

**Fig. 5.**
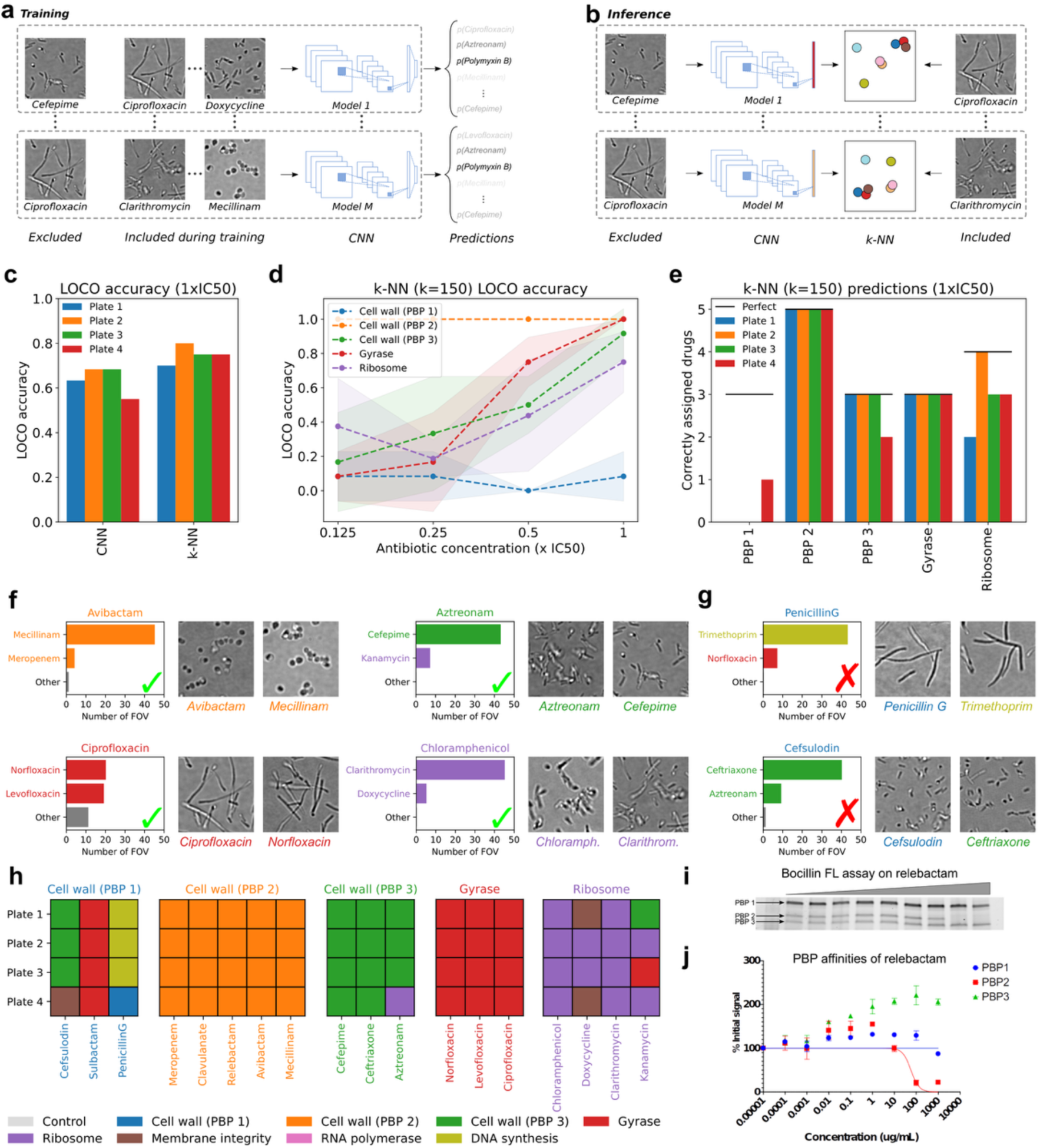
Assigning MoA of unseen compounds. **(a)** All images belonging to one antibiotic were systematically left out from the training data and models were trained in a cross-validated setting on three out of four plates. (**b**) The MoA of images belonging to the left-out antibiotic class was then determined via k-NN classification using the image embeddings of previously seen antibiotics on a hold-out test replicate. (**c**) Bars show the leave-one-compound-out (LOCO) MoA prediction accuracy at IC50 for each hold-out test plate, using either the CNN predictions (left: 63.8±5.4%) or the k-NN classifier (right: 75.0±3.5%). (**d**) The LOCO MoA accuracy was computed for images of bacteria exposed to different antibiotic concentrations with the k-NN classifier. The dashed lines show the average LOCO accuracy for each MoA and shaded areas indicate the standard deviation. (**e**) The number of correctly assigned antibiotics using k-NN classification is shown for all four hold-out test plates by MoA. The black horizontal lines indicate perfect assignment. (**f**) Examples of correctly assigned antibiotics showing the assignment of individual images to previously seen antibiotics with a k-NN classifier (k=150). Image crops of bacteria exposed to the left-out antibiotic and bacteria treated with antibiotics most commonly associated are shown. (**g**) Examples of incorrectly assigned antibiotics according to **Table 1** with image crops showing bacteria exposed to the left-out antibiotic and bacteria treated with the antibiotics most commonly associated. (**h**) LOCO predictions are shown for each hold-out test plate and antibiotics with colours indicating the assigned MoA. (**i,j**) The fluorescent penicillin analogue Bocillin FL was used in a competition assay against relebactam at increasing concentrations. (**i**) Gel showing bands of PBP1, 2 and 3. (**j**) Normalised fluorescence intensity (symbols and error bars show mean and standard deviation of three biological replicates) as function of relebactam concentration and fitted curves.

At the highest concentration (IC_50_), we observed good to excellent MoA prediction accuracy for compounds inhibiting PBP2 (100%) PBP3 (88.89%), DNA gyrase (100%) and ribosome (75%), while compounds targeting PBP1A/B were almost never correctly identified (1 out of 12 cases; **Fig. 5e**; **Discussion**).

We then investigated MoA assignments at the level of individual FoVs (**Fig. 5f**). For instance, in the case of avibactam, out of the 50 FoVs, 45 were classified as mecillinam and 4 as meropenem. Since both compounds target PBP2, avibactam was correctly classified as a PBP2 inhibitor. This classification is also consistent with the images, since both avibactam and mecillinam induce coccoid morphologies (**Fig. 5f**). Similarly, 20 FoVs of bacteria exposed to ciprofloxacin were classified as norfloxacin and 19 FoVs as levofloxacin which is consistent with the elongated phenotypes induced by these compounds. By contrast, MoA prediction failed for the three PBP1A/B inhibitors sulbactam, cefsulodin and penicillin G (**Fig. 5h**). Penicillin G was incorrectly associated with the DNA synthesis inhibitor trimethoprim, while cefsulodin was incorrectly associated with the PBP3 inhibitor ceftriaxone (**Fig. 5g**). While these predictions do not match our literature-based ground-truth MoA classes (**Table 1**), suggesting a limitation of our approach, they are in fact consistent with visual inspection of the images, which shows that the morphology of bacteria exposed to penicillin G, a PBP1A/B inhibitor, resembles that of bacteria treated with trimethoprim more so than that of bacteria treated with cefsulodin (the other PBP1A/B inhibitor that penicillin G should in theory be associated with) (**Fig. 5g**). A similar observation can be made about cefsulodin, a PBP1A/B inhibitor that produces morphologies more similar to those of the PBP3 inhibitor ceftriaxone than those of penicillin G (**Fig. 5g**). Overall, the predicted MoA based on targets reported in the literature (**Table 1**) was correct in 56 out of 72 cases (**Fig. 5h**).

As pointed out above, we initially labelled relebactam as a PBP2 inhibitor – despite a lack of evidence of its target in *E. coli –* based only on the drug’s structural similarity with avibactam. Our model indeed classified relebactam as a PBP2 inhibitor from the images alone (**Fig 5h**). To verify this prediction experimentally, we measured affinities of relebactam to multiple PBPs using a Bocillin FL competition assay (**Fig. 5i,j**) (**Materials and Methods**). This assay confirmed that relebactam indeed selectively inhibits PBP2 as predicted by our model.

The above-mentioned classification errors for PBP1A/B inhibitors arise either from limited performance of our model or misannotations of our ground truth data based on the literature (**Table 1**). To analyse this, we extended the Bocillin FL competition assays (**Materials and Methods**) to determine the specificity against different PBPs of a subset of PBP inhibitors studied here, namely clavulanate, cefepime, penicillin G, cefsulodin and sulbactam (**Supp.** Fig. 8). Our data confirm that clavulanate, classified by our LOCO analysis as a PBP2 inhibitor (**Fig. 5h**), is indeed highly specific to PBP2, in agreement with our literature-based annotations (PBP1A/B: 256.9 μg/mL, PBP2: 0.615 μg/mL, PBP3: 1,047.0 μg/mL). Similarly, these data show that cefepime primarily inhibited PBP3, in agreement with our model predictions and annotation (PBP1A/B: 1.724 μg/mL, PBP2: 0.067 μg/mL, PBP3: 0.001 μg/mL). However, the affinity data for two of the three drugs we had labelled as PBP1A/B inhibitors were not consistent with the literature. Indeed, whereas our assay confirmed that cefsulodin primarily targets PBP1A/B (PBP1A/B: 0.748 μg/mL, PBP2: >10,000 μg/mL, PBP3: 10.4 μg/mL), in agreement with the literature annotation, neither penicillin G (PBP1A/B: 7.597 μg/mL, PBP2: 64.64 μg/mL, PBP3: 1.977 μg/mL) nor sulbactam (PBP1A/B: 37.4 μg/mL, PBP2: 200.3 μg/mL, PBP3: 102.2 μg/mL) had a specific target, but instead acted as pan-PBP inhibitors. Although these data do not explain the model predictions for cefsulodin, the lack of specificity of penicillin G and sulbactam is a likely cause of their inconsistent assignments by our model (see **Discussion**).

Thus, our data indicate that our CNN can learn to extract embeddings from BF images alone that enable accurate MoA prediction of an unseen drug if the MoA is represented by reference compounds with consistent target specificity.

### Detecting drugs with unseen modes of action

While the above results indicate that known MoAs can be assigned to novel compounds, addressing AMR requires the ability to identify compounds with novel MoAs. To address this, we assessed if our approach is capable of detecting compounds with a MoA that - unlike in our previous analysis - was not seen by our model during training. We designed a leave-one-MoA-out (LOMO) experiment where we singled out one MoA and trained our CNN on BF images of drug-treated bacteria after systematically leaving out all compounds from this MoA (**Fig. 6a**). We then computed outlier scores on image embeddings with a local outlier factor (LOF) algorithm (16) (**Fig. 6b**). To serve as negative controls for novel MoA detection, we randomly removed one compound for each MoA with at least three compounds (**Table 1**) from the training data.

**Fig. 6.**
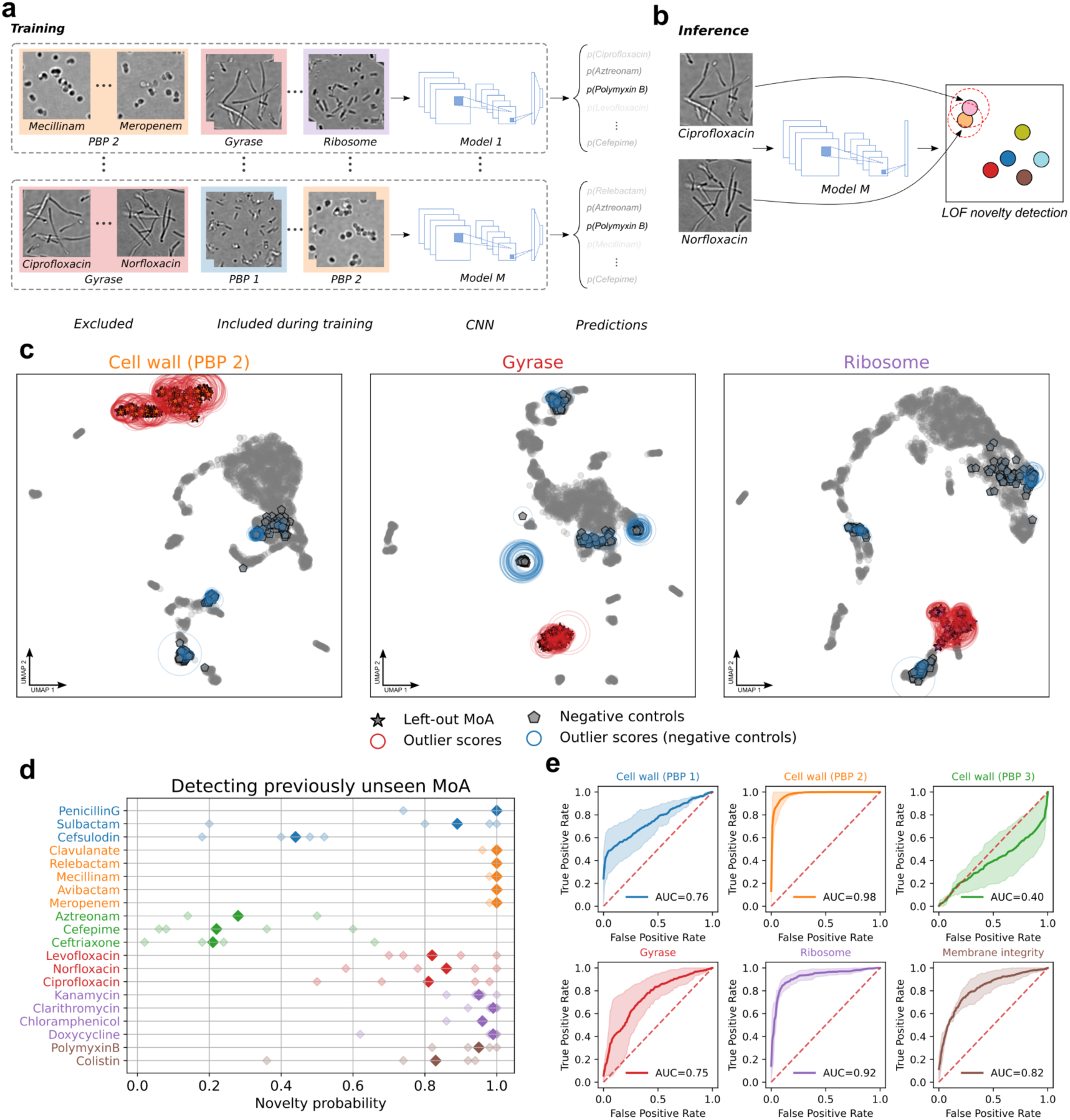
Detecting antibiotics with novel MoAs. **(a)** All images belonging to the same MoA were systematically left out from the training data and models were trained in a cross-validated setting on three out of four plates. (**b**) Embeddings from images of the left-out MoA were then obtained and outlier scores assigned with a LOF algorithm. (**c**) UMAPs of embeddings for models trained on compounds from all MoAs except PBP2 (left), DNA gyrase (centre) and ribosome (right). Unsupervised outlier detection was conducted using LOF on embeddings of bacteria exposed to all concentrations (grey dots). Embeddings of compounds from the held-out MoA are shown as stars, negative controls as pentagons and their outlier scores as red and blue circles respectively. (**d**) For each compound, probability scores of MoA novelty were computed by thresholding outlier scores and computing the fraction of detected embeddings (one for each MoA). Pale symbols show the novelty detection probability for each plate and the median is shown in bright colours. Colours indicate the MoA. (**f**) Cross-validated receiver operating characteristic (ROC) curves were obtained for each held-out MoA on four hold-out test plates with mean AUC values shown.

After model training, we obtained embeddings from images of bacteria on a hold-out test plate treated with all compounds except those of the left-out MoA and negative controls at all four concentrations (grey dots in **Fig. 6c**). Next, we defined an image as showing a novel MoA if the outlier score of its embedding exceeded a threshold (and as a known MoA if it did not). We then computed “novelty probabilities” for each left-out compound (at the IC_50_) by considering the fraction of FoVs that were detected as exhibiting a novel MoA based on their embeddings (**Fig. 6d**). The compounds of the left-out MoA and the negative controls allowed us to quantify true and false positives, respectively. We determined ROC curves by cross-validation across four hold-out test plates and computed mean AUC values for each MoA (**Fig. 6e**).

All compounds targeting PBP2, DNA gyrase, ribosome and membrane integrity exhibited median novelty probabilities above 0.8 and high or very high AUC scores (PBP2: 0.98, DNA gyrase: 0.75, ribosome: 0.92, membrane integrity: 0.82), indicating that these compounds with these MoAs induce phenotypes that can be well differentiated from the MoAs seen by the model during training. In the case of PBP1A/B (AUC=0.76), only penicillin G and sulbactam exhibited high novelty probabilities, whereas cefsulodin did not (**Fig. 6d**). For the DNA gyrase inhibitors, median novelty scores across compounds were consistently high (AUC=0.75), but individual scores by plate exhibited a higher degree of spreading compared to Ribosome or PBP2 inhibitors. Conversely, PBP3 inhibitors consistently showed low novelty probabilities (AUC=0.40), suggesting that these compounds induce phenotypes that do not differ from other MoAs sufficiently strongly to be determined as novel MoAs *de novo*. Note that the PBP1 inhibitor cefsulodin was predominantly classified as a PBP3 inhibitor in our LOCO analysis (**Fig. 5h**) indicating that cefsulodin induces PBP3-like morphologies. The consistently low novelty scores observed for PBP3 inhibitors could thus explain the low novelty scores observed for cefsulodin.

Taken together, our results show that compounds from all but one of the studied MoAs induce phenotypes that are robustly detected as novel when excluded from the training data. Our approach could thus play a key role in identifying promising compounds with potentially novel MoAs in phenotypic screens.

## Discussion

In this study, we introduced a deep learning (DL) approach based on convolutional neural networks (CNNs) that can extract meaningful phenotypic features from microscopy images of bacteria treated with antibiotic compounds and showed that these features can be used for multiple downstream tasks with high relevance for phenotypic drug screening. These tasks include the recognition of MoA or even compound identity directly from images (**Fig. 2**), the detection of weak phenotypes at subinhibitory drug concentrations (**Fig. 4**), the assignment of drug MoAs for novel compounds (**Fig. 5**) and the identification of compounds with novel MoAs (**Fig. 6**). Our work builds on prior studies in eukaryotic cells and bacteria that established cytological profiling to analyse drug-induced phenotypes and group drugs by MoA, whether using handcrafted features (7,8), or more recently, DL approaches (11). However, our study represents a significant extension of these early approaches in several ways.

First, we established that DL can extract both MoA- and compound-specific features from brightfield (BF) images alone, enabling perfect MoA recognition without any fluorescent labelling (**Fig. 3c,e**). This came to us as a surprise, since most phenotypic drug screening assays employ a mixture of fluorescent dyes to characterise subcellular features, as illustrated by the popular Cell Painting assay (17–19) or bacterial cell profiling assays (5,7,8,11). Our finding that fluorescent labelling of bacterial membranes and DNA is redundant with BF images is consistent with studies showing that DL models can be trained to generate realistic fluorescent labelling or coloration (virtual staining) *in silico* from label-free images in eukaryotes (20) and in bacteria respectively, as illustrated e.g. by a recent report that Gram-staining can be predicted from darkfield images of *E. coli* and *Listeria innocua* (21). Our results extend previous work in eukaryotic cells showing that DL applied to BF images performs on par with classical CellProfiler features to determine drug MoA (12) to the case of bacteria. More importantly, our findings enable a drastic simplification of the experimental workflow of phenotypic imaging screens by removing the need for staining and protocol optimisation, as well as avoiding the associated costs and potential confounding factors like variations in fluorescent staining. The removal of staining also opens the door to phenotypic screening in live cells, potentially circumventing fixation issues and enabling the incorporation of live cell dynamics in phenotyping.

Second, although DL-based image recognition for natural images typically requires thousands of training images per class to reach state-of-the-art classification accuracies, we showed that on the order of seven FoVs per condition (i.e. compound and concentration) are sufficient to achieve high accuracies in MoA recognition and that acquiring more images does not yield significant improvements (**Fig. 3g-i**). The main reason for this surprisingly low number is probably the large number of cells contained in each FoV, and the relative simplicity of *E. coli* BF images compared to natural images like those present in the ImageNet dataset. Nevertheless, our observation that pretraining our model on ImageNet significantly reduces the required number of images suggests that many ImageNet features are also relevant for BF *E. coli* images. Our finding that recognition accuracy saturates with only a handful of FoVs should encourage future phenotypic imaging screens to reduce image acquisition time per well and thereby increase the rate of assayed compounds.

Third, we emphasise that our model was trained to predict the combinations of drugs and concentrations and thus has no explicit prior knowledge of the MoA. Despite this, we show that our model implicitly encodes similarities by MoA in its latent space. This is a welcome feature, as it means that a straightforward supervised learning strategy with well-defined and unambiguous labels (drugs and concentrations) can learn biologically meaningful representations from the image data alone.

Fourth, we have shown that DL can robustly detect weak phenotypes induced by drugs at concentrations well below the inhibitory concentrations (detection AUC=0.81±0.08 at 0.125×IC_50_). This result implies that large chemical libraries at a fixed concentration can be screened with high sensitivity, i.e. with a high likelihood of picking up active compounds that elicit subtle phenotypes in the images without affecting bacterial growth. In a second step, these hit compounds could then be titrated to determine their IC_50_ values, imaged at IC_50_, and analysed for MoA assignment with high accuracy at IC_50_ concentrations for most MoAs (PBP2: 100%, PBP3: 91.7%, DNA gyrase: 100%, ribosome: 75%) except PBP1A/B (8.3%, **Fig. 2c, 5e**).

Finally, we have addressed the critical task of determining if a novel compound also has a novel MoA. We have shown robust novelty detection for four out of five tested MoAs, with excellent performance (specificity and sensitivity) for PBP2 and ribosome inhibitors (AUC=0.98 and 0.92, respectively), good to very good performance for inhibitors of PBP1A/B, DNA gyrase and membrane integrity (AUC=0.76, 0.75, and 0.82, respectively), but acknowledge poor performance for PBP3 inhibitors (AUC=0.40). Despite imperfections, our novelty detection opens the door to screening large libraries of compounds using high-throughput microscopy to identify compounds that are likely to act through novel MoAs. We acknowledge several limitations or open questions. While MoA assignment in the LOCO analysis was good for compounds belonging to four MoA groups, it is notable that none of the compounds labelled as PBP1A/B inhibitors were correctly assigned. However, inspecting images of *E. coli* treated with different PBP1A/B inhibitors (**Supp.** Fig. 2a) reveals that these misclassifications are likely due to inconsistent phenotypes induced by these compounds (**Fig. 5g**). For instance, bacteria treated with penicillin G exhibit long filamentous morphologies similar to those induced by trimethoprim, while bacteria treated with cefsulodin exhibit shorter filaments and are thus more similar to morphologies observed for PBP3 inhibitors. These phenotypic inconsistencies and the results of our PBP affinity assays call into question the grouping of these PBP1A/B inhibitors based on the literature (**Table 1**). Cefsulodin has been reported to selectively target PBP1A/B (PBP1A: 0.47 μg/mL,PBP1B: 3.7 μg/mL,PBP2: >250 μg/mL and PBP3: >250 μg/mL) (22). However, this is not supported by our PBP affinity assay (**Supp.** Fig. 8), where we observed inhibition of both PBP1A/B (0.75 μg/mL) and PBP3 (10.4 μg/mL). Sulbactam has been reported to primarily target PBP1A with low affinities for PBP2 and PBP3 (24), but this is also not in line with our affinity assays, where we observe broad inhibition of various PBPs (**Supp.** Fig. 8). By contrast, the high reported affinity of penicillin G for multiple PBPs (PBP1A: 0.5 μg/mL, PBP1B: 3.0 μg/mL, PBP2: 0.8 μg/mL, PBP3: 0.9 μg/mL and PBP4: 1.0 μg/mL) (23) is consistent with our PBP affinity assay showing a pan-PBP inhibitor activity (**Supp.** Fig. 8). Taken together, these differences in PBP affinities between putative PBP1A/B inhibitors plausibly explain the observed inconsistencies in bacterial phenotypes. We note that the differences in PBP affinities between our experiments and prior studies could result from variations in bacterial strains due to their individual history of pre-exposure to β-lactams. Cefsulodin is the only drug in our dataset that primarily targets PBP1A/B (0.75 μg/mL), although with high secondary affinity for PBP3 (10.4 μg/mL). Nevertheless, it is classified as a PBP3 inhibitor by our LOCO analysis (**Fig. 5g**). We note that recent work has shown that PBP1A and PBP1B do not regulate the morphology of *E. coli* bacteria (25) and instead contribute to cell-wall integrity, whereas PBP3 directly regulates bacterial cell shape. If inhibition of PBP1A/B does not produce a visible phenotype, it appears likely that morphological changes will be driven by inhibition of PBP3 alone. This would be consistent with our model’s classification of cefsulodin as a PBP3 inhibitor.

Additionally, our study included four β-lactamase inhibitors (sulbactam, avibactam, clavulanate and relebactam). Although, sulbactam has been reported to preferentially bind PBP1A, we show it is a pan-PBP inhibitor (**Supp.** Fig. 8), while we confirm that avibactam and clavulanate target PBP2 as reported. Relebactam, however, has not been studied in *E. coli* and our LOCO analysis indicates that this drug induces PBP2-like phenotypes similar to avibactam and clavulanate (**Fig. 5h**). Indeed, our affinity assay confirmed that relebactam selectively targets PBP2 in our *E. coli* strain (**Fig. 5g**). This is consistent with data on other DBOs like avibactam that have PBP2 as their target. Interestingly, prior work shows that avibactam preferentially binds PBP1B and PBP4 in *Pseudomonas aeruginosa* while relebactam acts as a pan-PBP inhibitor (25), although their target predictor classifies both as PBP2 inhibitors consistent with our data.

Intriguingly, while trimethoprim is commonly classified as a DNA synthesis inhibitor and specifically targets folate synthesis, some studies have shown that trimethoprim also directly targets the peptidoglycan pathway (26,27). Indeed, peptidoglycan synthesis relies on nucleotide-activated sugars such as UDP-N-acetylglucosamine and UDP-N-acetylmuramic acid. Inhibition of nucleotide synthesis by trimethoprim would thus also affect peptidoglycan synthesis and more generally cell wall biogenesis that relies on many nucleotide-activated sugars including Lipopolysaccharides. The assignment of penicillin G to trimethoprim appears consistent with these prior findings.

Taken together, these observations highlight the limitation of grouping compounds into discrete MoA classes and using reference compounds that inhibit multiple targets. One promising avenue to overcoming this limitation is to predict drug targets by comparing their induced phenotypes to those of large mutant libraries. As another caveat, our results are limited to *E. coli* bacteria, which are relatively large and exhibit strong morphological changes in response to drug exposure. Future work should thus explore if our BF-only methodology extends to other bacterial species with more subtle morphological phenotypes such as *Staphylococcus aureus* or *Acinetobacter baumannii*.

Overall, our work is a step towards deploying state-of-the-art computational analysis methods, notably DL-based analysis of microscopy images, to bacterial phenotypic screens. We propose that our CNN approach and the downstream analysis methods described here could form the basis for a comprehensive phenotypic screening pipeline to discover novel classes of antibiotics to fight AMR.

## Methods

### Bacterial growth protocol for *E. coli*

*E. coli* K-12 MG1655 was grown in 5 mL of cation adjusted Mueller-Hinton Broth II (MHBII, Fisher Scientific cat. 11800482) overnight at 37°C until stationary phase. The culture was then diluted to 0.05 OD in fresh MHBII and grown at 37°C for 1h 45min until mid-exponential phase between 0.5 to 0.7 OD before further dilution. IC_50_ values were determined via broth serial dilutions in flat-bottomed 96-well plates (TPP cat. 92096) with a starting inoculum density of 0.05 OD in a total volume 100μL. Micro dilution plates were incubated at 37°C, 180 rpm for 18 h. The final cell density was measured on a Tecan Spark Multimode Plate Reader and plotted on GraphPad Prism to calculate the IC_50_ values.

For microscopy, antibiotics (see **Supp. Table 2** for further information on the source of the antibiotics) were dispensed into 96-well plates in nanoliter volumes using an Echo 550 liquid handler (Beckman Coulter) before adding 100 µL of bacteria from mid-exponential growth diluted to 0.05 OD. The plates were incubated at 37°C, 180 rpm for 2h 15 min. Then, 20 μg/mL of FM4-64FX (Invitrogen cat. F34653) was added and was further incubated at 37°C, 180 rpm for another 15 min. The supernatant was then removed after centrifugation at 4000 x g for 5 min. 100 µL of fixative (2.8% paraformaldehyde, 0.04% glutaraldehyde in PBS) was added and the plates were incubated at room temperature at 800 rpm for 30 min. Cells were washed twice with 200 µL of PBS by centrifugation and resuspension. Then, cells were suspended in 100 µL of PBS containing 10 µg/mL of Hoechst 33342 (Invitrogen cat. H1399) and transferred to PhenoPlates (Revvity cat. 6055300) to determine the OD at 600 nm. Finally, the entire plate was diluted based on the median OD of the untreated wells to a calculated OD of 0.036 in 100µL. The fixed and diluted cells were stored at 4°C until imaging on the Opera Phenix Plus (Revvity).

### Penicillin-binding protein affinity assay

E. coli K-12 MG1655 was grown in 5 mL of LB overnight at 37°C until stationary phase. The culture was then diluted to 0.05 OD in fresh LB and grown at 37°C for 1h 30min until mid-exponential phase between 0.5 to 0.7 OD. Bacteria were collected by centrifugation at 3220 x g for 5 min and washed with 50 mM Tris, pH 7.8. Then, bacteria were suspended in 50mM Tris, 200mM EDTA, pH 7.8 to 20 OD mL. 2 5μL of 10-fold serially diluted antibiotics starting at 1 mg/mL were aliquoted into a V-bottomed plate and 25 μL of 20 OD bacteria was added. The plate was incubated at 20°C for 30 minutes and then centrifuged at 3220 x g for 5 minutes to pellet the cells. The supernatant was removed, and cell pellets were suspended in 50 μL of 50 mM Tris, 200 mM EDTA, pH 7.8 containing 25 μM of Bocillin-FL (Invitrogen cat. B13233). The plate was incubated at 20°C for 30 minutes, 1400 rpm. The cells were collected by centrifugation, suspended in 100 μL of 50 mM Tris pH 7.8, and centrifuged again before removing the supernatant. 50 μL of Laemmli sample buffer was added and the plate was incubated at 70°C for 10 minutes at 800 rpm before centrifugation at 3220 x g for 10 minutes to pellet insoluble materials. 25 μL of each sample was run on 4-15% Tris-Glycine SDS-PAGE (Bio-Rad, cat. 4568084). The gels were imaged on a Bio-Rad Chemidoc at 488 nm to visualize the Bocillin-FL labelled bands and at stain-free settings to visualize total protein content.

### High-throughput imaging

Prior to imaging, plates stored at 4°C were left for at least 20 min at room temperature and then centrifuged at 35 rcf for 10 min to bring floating bacteria to the well bottom. Images were then acquired using an automated Opera Phenix Plus High-Content Screening System (Revvity) with 63×/NA 1.15 water immersion objective in widefield mode without binning. Two imaging sequences were set up: one for the simultaneous acquisition of fluorescent markers through two different cameras, and one for the BF acquisition via a dedicated camera. Hoechst 33342 and FM4-64X were imaged with a 405 nm or 488 nm excitation laser lines and a 435-480 nm or 650-680 nm bandpass filters, respectively. Images were acquired with exposure times of 80 ms for BF, 400 ms for FM4-64FX and 250 ms for Hoechst 33342. In total, 50 FoVs were acquired in non-overlapping randomly chosen positions of the whole area of each well. These random positions in each well were the same on individual plates but varied between plates.

### Image preprocessing

Images for training were obtained by converting high-throughput images from 16-bit to 8-bit and clipping pixel intensities by percentile between 0.1 and 99.9 to remove spurious saturated pixels. Images from three out of four plates were then randomly split into training (80%) and validation (20%) set. During training, full-resolution images with 2,160×2,160 pixels were passed through a data augmentation pipeline: images were randomly flipped, rotated and random changes in brightness, contrast, saturation and hue (for images with more than one channel) were applied. To account for potential edge effects due to random rotation, full images were further centre cropped (1,528×1,528 pixels). Next, from this centre-crop, a random crop of size 1,500×1,500 pixels was obtained and images subdivided into nine tiles of size 500×500 pixels. Tiles were then resized to 256×256 pixels and pixel intensities normalised between -1 and 1.

### Deep learning model training

The DL model was trained for 250 epochs using an Adam (28) optimiser with a learning rate of 0.001. A batch-size of 16 images was used. Note that the effective batch-size was 144 (16×9), since each image was further subdivided into nine tiles. During training, tiles were passed through an EfficientNet-B0 backbone, with weights initialised by pretraining on ImageNet^1^ [^1^https://download.pytorch.org/models/efficientnet_b0_rwightman-7f5810bc.pth] (except where indicated otherwise), and latent feature vectors corresponding to individual tiles aggregated by averaging to obtain a global feature vector for the full-resolution input image. The model was trained to predict the combination of antibiotic treatment and concentration as a multi-class classification task with a cross-entropy loss using L2 regularisation with λ=0.001 to avoid overfitting. Models were trained with PyTorch 2.3.1 (29) in Python 3.9 on the HPC cluster at Institut Pasteur Paris. See **Supp. File 1** for the full model architecture.

### Estimating minimum number of required images

To estimate the minimum number of required training images, we fit an exponential *f*(*x*) = *a* (1 −*e*^−*bx*^) (where *x* is the number of images per condition and *f*(*x*) the hold-out test accuracy) to mean MoA accuracies obtained from hold-out test data with models trained on different numbers of images per well ranging from 3 to 120. From these exponential fits, the plateau *a* was obtained and the number of images per well estimated to achieve 95% of this MoA classification accuracy plateau. We used the curve fitting function from the scipy package in Python 3.9 (30).

### MoA assignment with k-NN classifier

Embeddings from images of previously seen conditions were obtained from a hold-out test plate. A k-NN classifier was then applied to these embeddings corresponding to the highest concentration (IC_50_) using MoA labels as targets with the scikit-learn package (31) in Python 3.9. The number of k neighbours was set to 150 using cosine distance (1 minus the cosine similarity) as the metric and weighing points by the inverse of their distance thereby ensuring that closer neighbours had a greater influence on the MoA assignment than neighbours which were further away. To predict the MoA, embeddings of previously unseen compounds were obtained from the same hold-out test plate. This analysis was repeated for each model with a left-out compound across four hold-out test plates.

### Novelty detection with local outlier factor algorithm

Embeddings from images of previously seen conditions were obtained from a hold-out test plate. A LOF classifier was applied to these embeddings using all available concentrations with the scikit-learn package in Python 3.9. The number of k neighbours was set to k=9 and cosine distance (1 minus the cosine similarity) was used as the distance metric. To determine if a compound was an outlier, i.e. had a novel MoA, the local outlier factor on embeddings from unseen conditions at the highest concentration (IC_50_) was computed and a threshold applied as described in (16). For each compound, novel MoA detection probabilities were computed as the fraction of detected outliers considering all FoVs. ROC curves were obtained by computing the local outlier factor for previously unseen compounds and embeddings corresponding to left-out negative control conditions (one per MoA with at least three compounds) and positive controls (i.e. compounds with the left-out MoA). AUC values were computed with the scikit-learn package. This analysis was repeated for each model with a left-out MoA across four hold-out test plates.

## Acknowledgements

We thank C. Thépenier, C. Spahn, S. Jang, D. Shum and N. Aulner for insightful comments on the manuscript. We thank A. Zettor for preparing plates with an acoustic liquid handler at the Chemogenomic and Biological Screening Platform (PF-CCB) at Institut Pasteur. D.K. was funded by the Pasteur-Paris University International doctoral program (PPU), the INCEPTION program (Investissement d’Avenir grant ANR-16-CONV-0005) and a Fondation pour la Recherche Médicale (FRM) Fin de Thèse (FDT) grant. K.K. was supported by the INCEPTION program, the Pasteur-Roux-Cantarini Postdoctoral Fellowship (Institut Pasteur) and the Fondation pour la Recherche Médicale (FRM) grant EQU202403018034. J.P. was funded by Fondation pour la Recherche Médicale (grant number EQU202303016284 to Pedro Alzari). Work in the Boneca lab was supported by the Fondation pour la Recherche Médicale (FRM) grant EQU202403018034. The UTechS Photonic BioImaging, C2RT, Institut Pasteur, is supported by the French National Research Agency (France BioImaging, ANR-24-INBS-0005 FBI (BIOGEN); Investments for the Future). We acknowledge Institut Pasteur and the Région Île-de-France (DIM1Health program) for the use of the Opera Phenix system. We also acknowledge the INCEPTION program for funding a GPU farm used in this work.

