## Supplementary figures and images for "Deep learning recognises antibiotic modes of action from brightfield images"

### Supplementary File 1

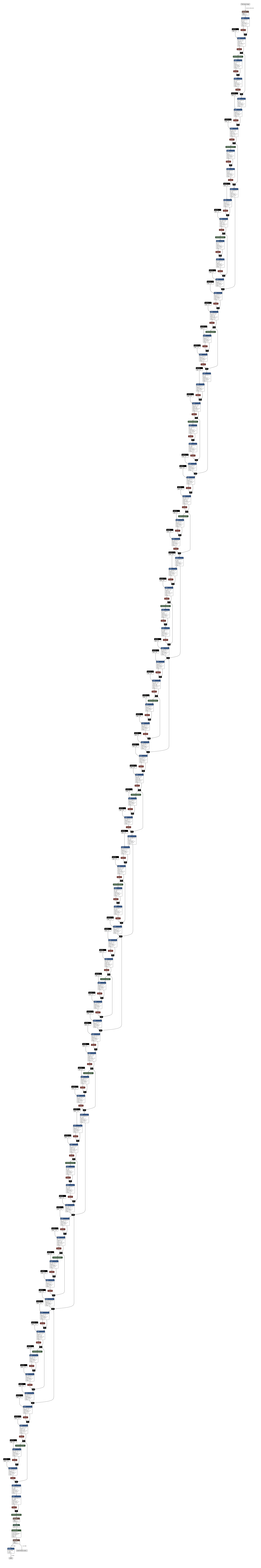

### Supplementary File 2

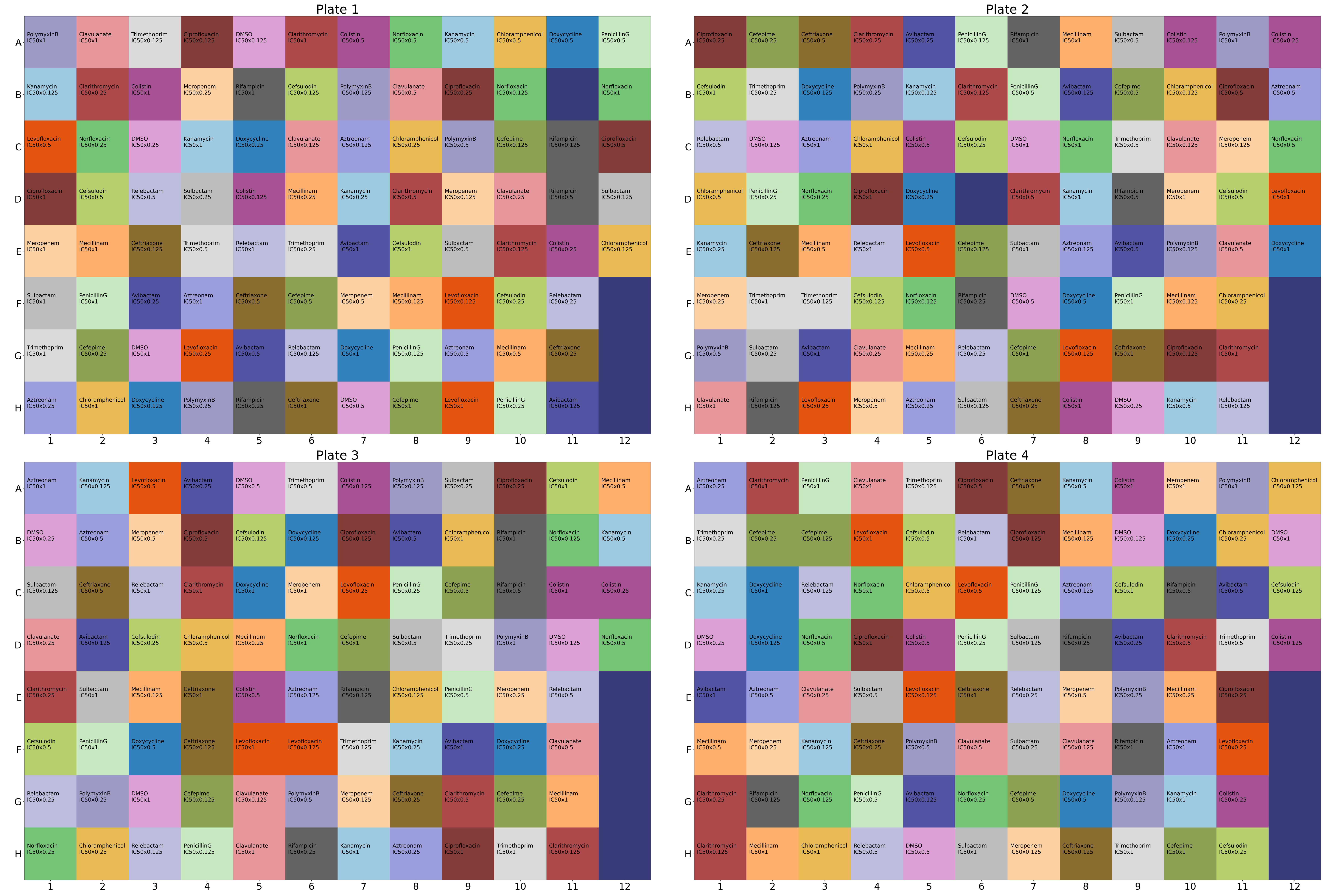
