## Supplementary Material for "Deep learning recognises antibiotic modes of action from brightfield images"

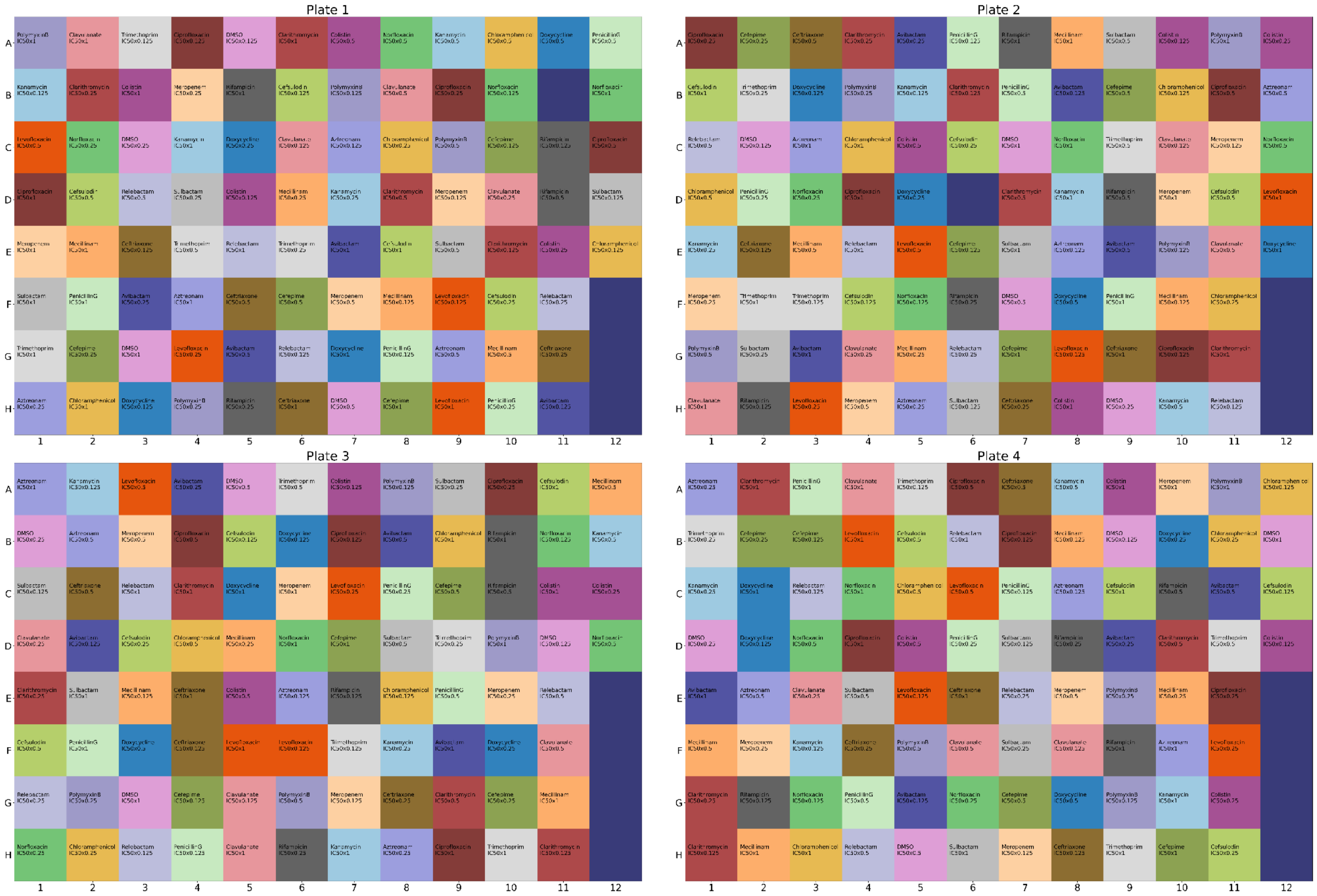

**Supp Fig. 1: Pseudo-randomised plate layouts.**
Each plate was acquired using independently cultured *E. coli* bacteria (biological replicate) and pseudo-block-randomised plate layouts[1] were generated for compound distribution with an acoustic liquid handler at four different concentrations. Pseudo-randomisation was used to decorrelate the treatment conditions from the positions on the plate and alleviate potential confounding effects. Colours in the plate maps correspond to individual antibiotics. See **Supp. File 2** for a high-resolution image.

[1] https://github.com/crukci-bioinformatics/PlateLayout

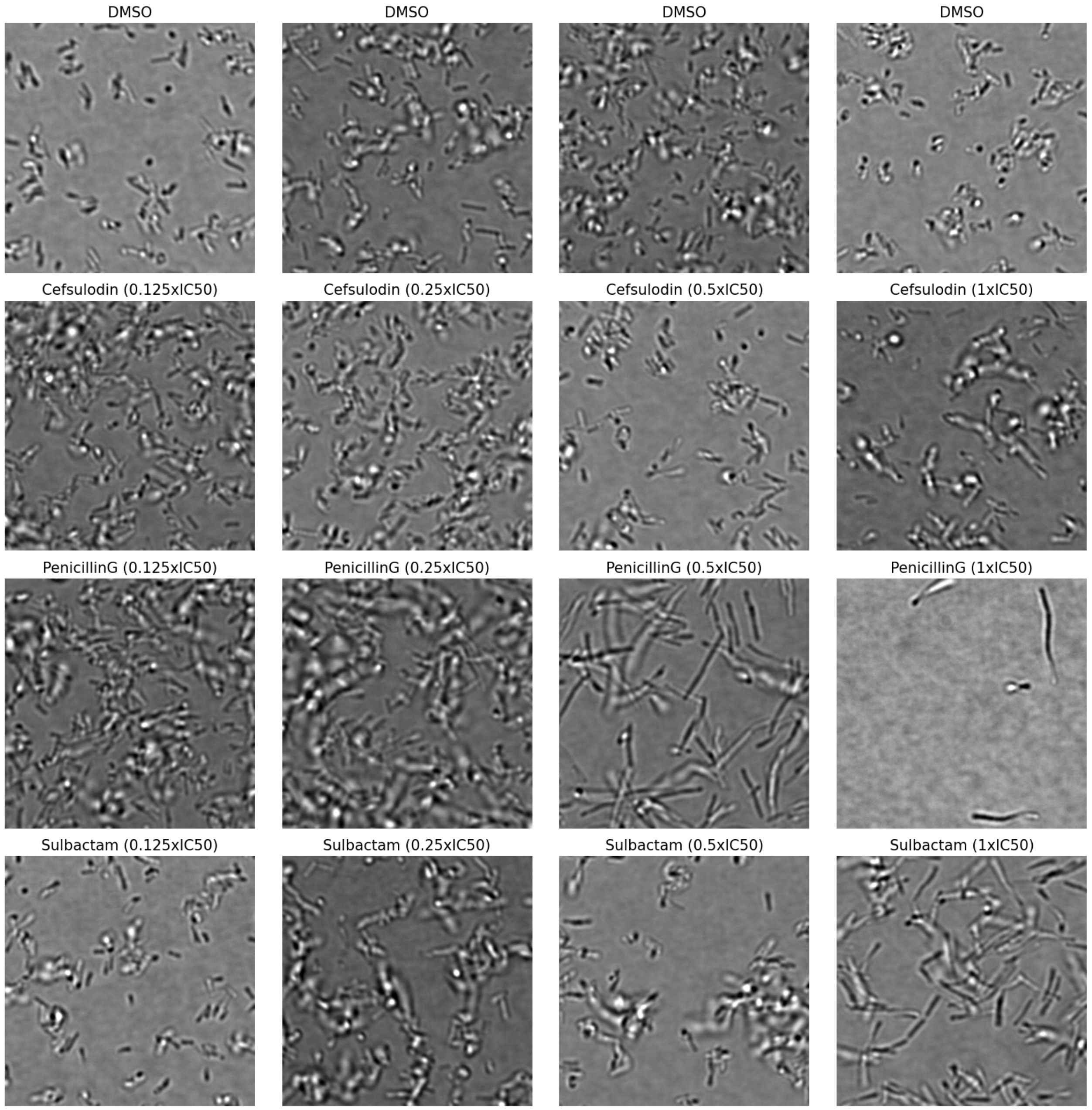

**Supp Fig. 2a: Image gallery for PBP1A/B inhibitors.**

In Supp. Figs. 2a-f, randomly chosen and centre-cropped (1,000×1,000 pixels) BF images are shown for bacteria treated with all compounds at all concentrations and grouped by MoA: PBP1A/B (**a**), PBP2 (**b**), PBP3 (**c**), gyrase (**d**), ribosome (**e**), membrane integrity, RNA polymerase and DNA synthesis (**f**).

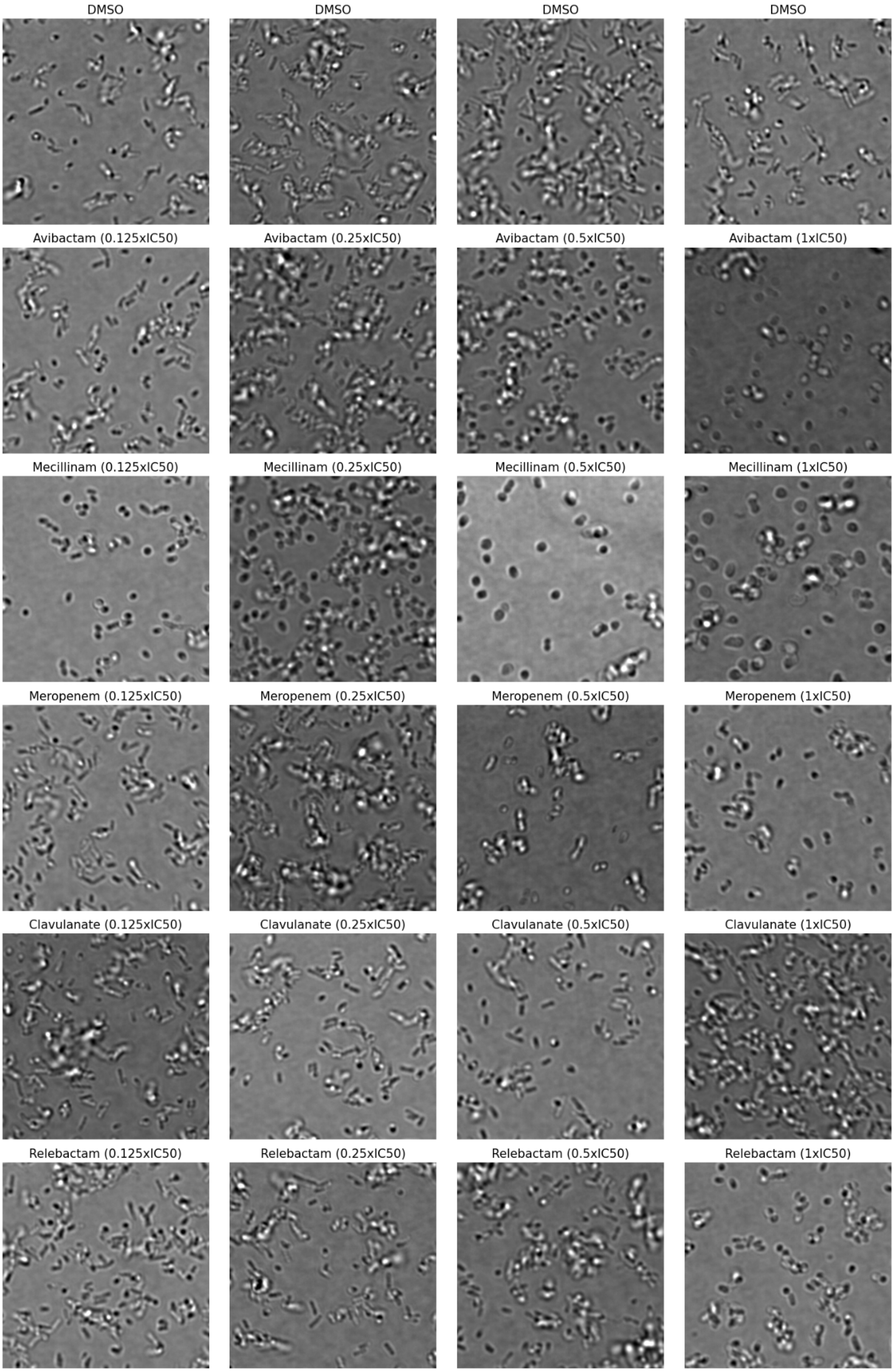

**Supp Fig. 2b: Image gallery for PBP2 inhibitors.**

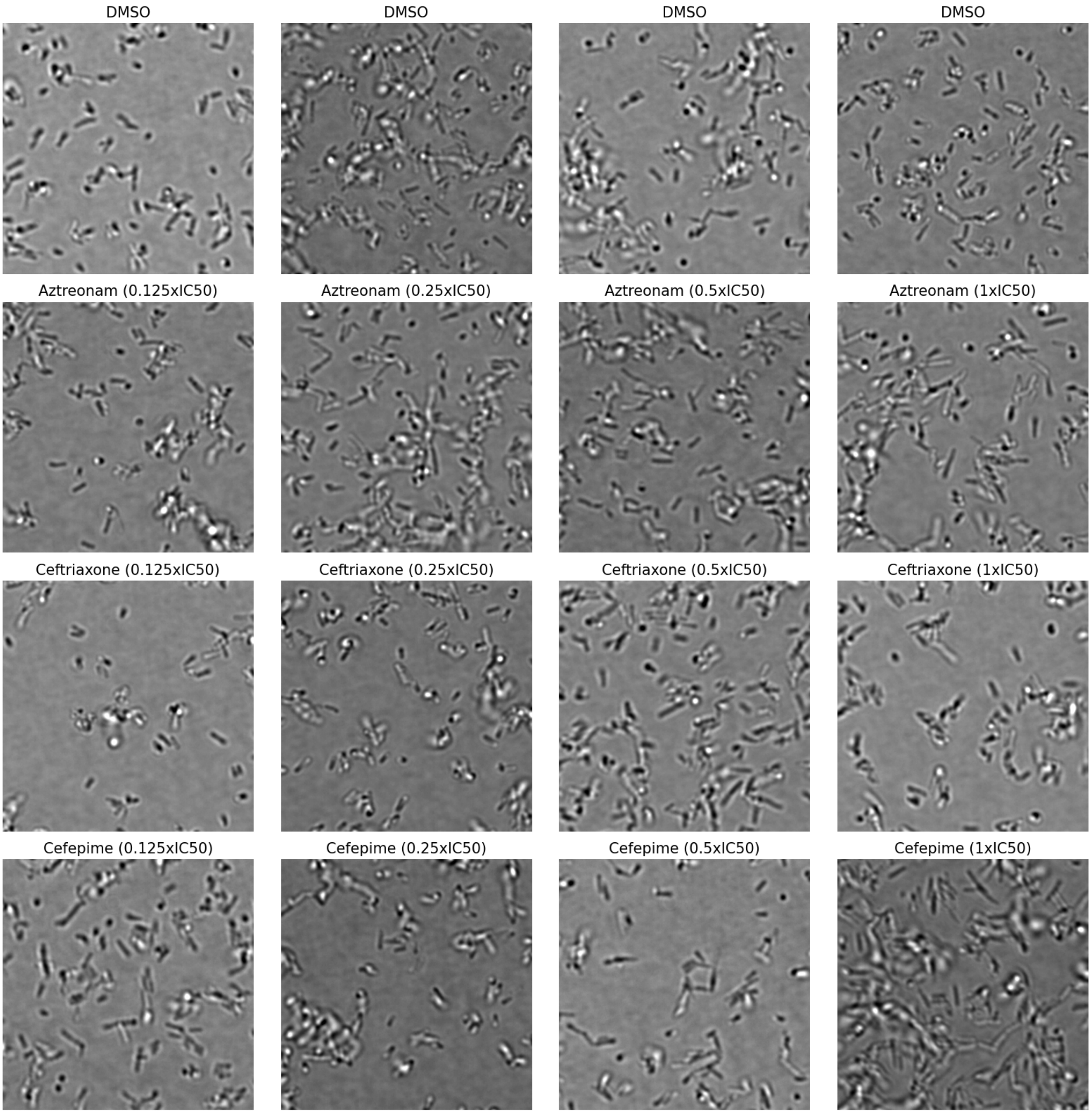

**Supp Fig. 2c: Image gallery for PBP3 inhibitors.**

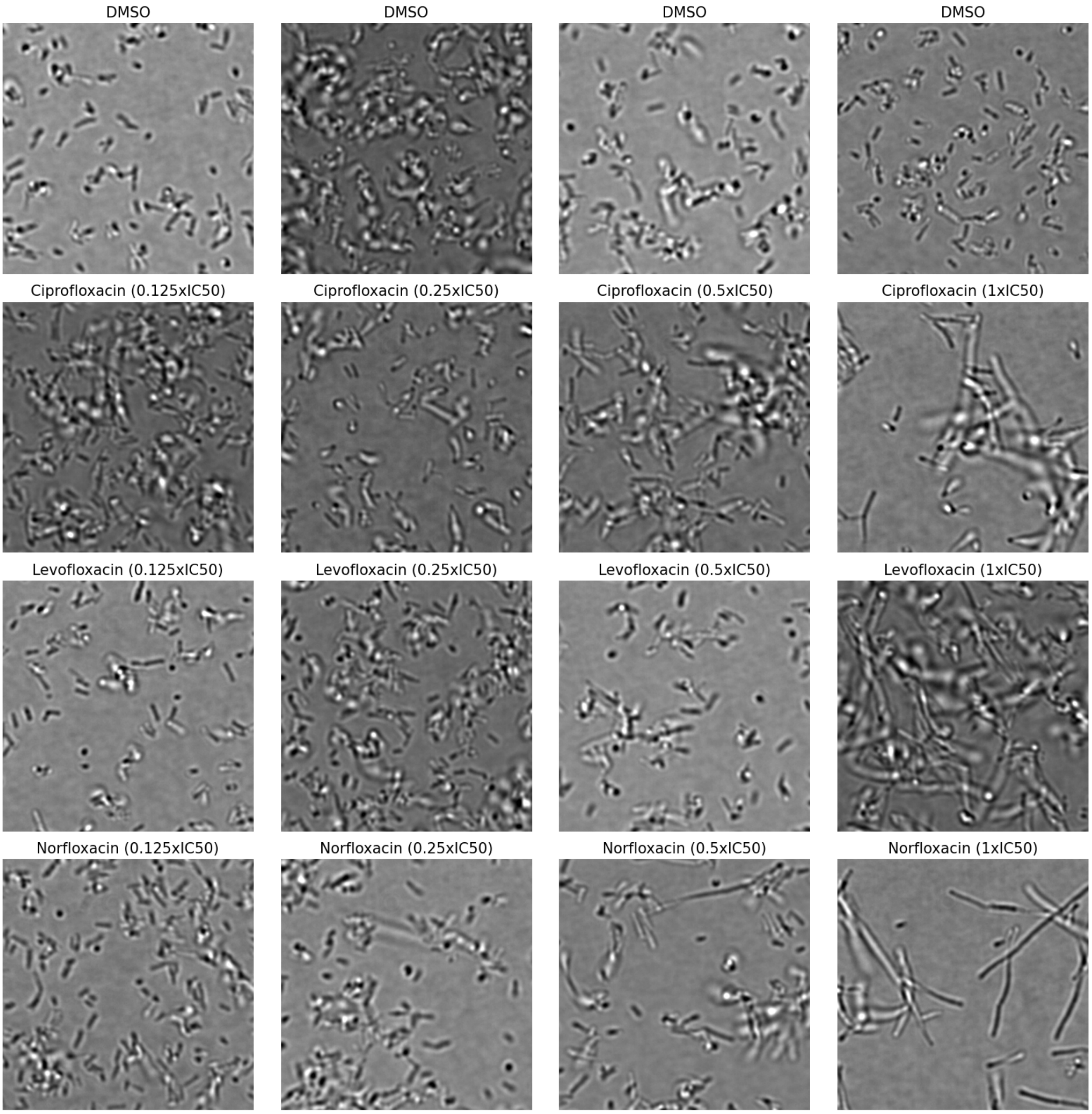

**Supp Fig. 2d: Image gallery for DNA gyrase inhibitors.**

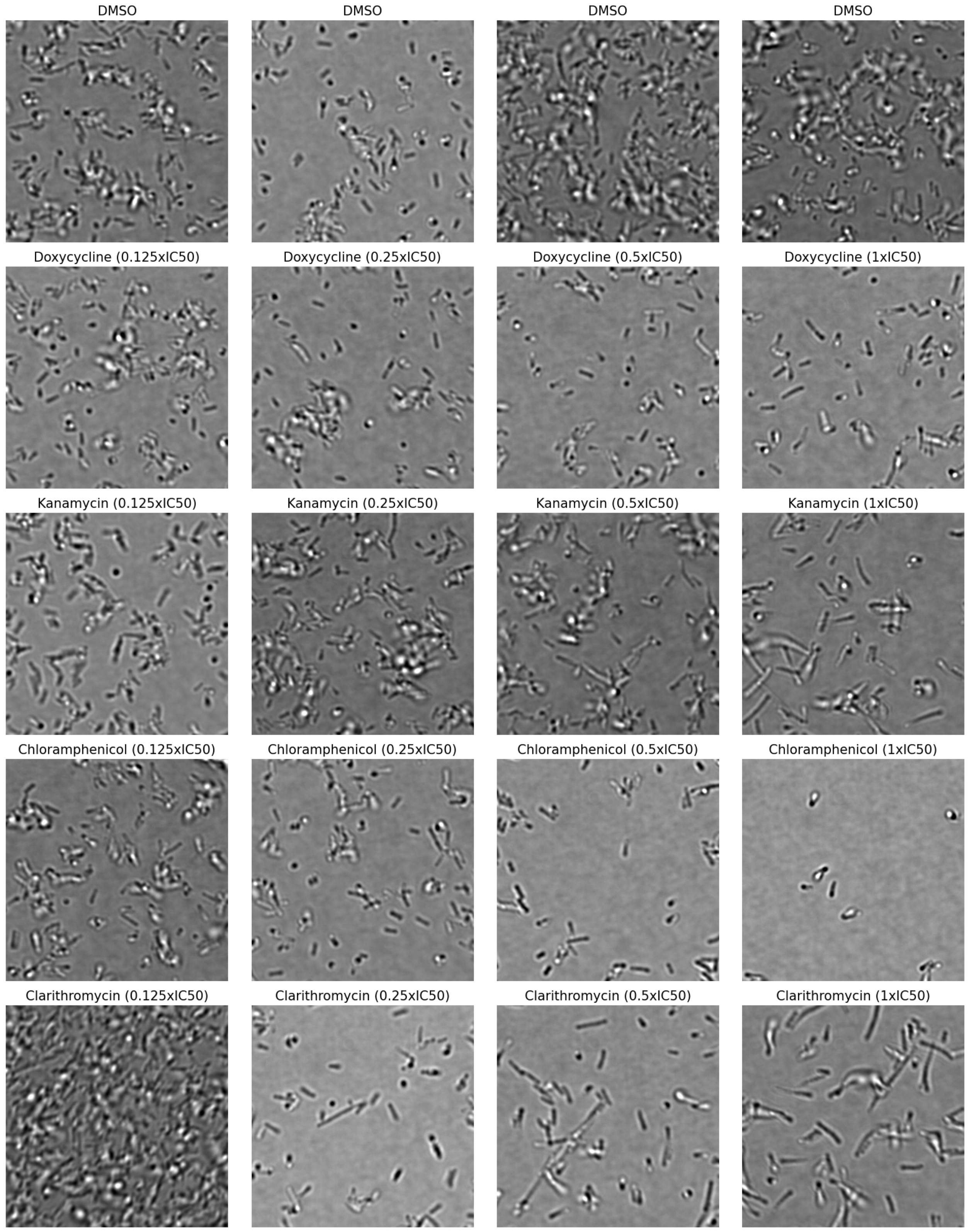

**Supp Fig. 2e: Image gallery for ribosome inhibitors.**

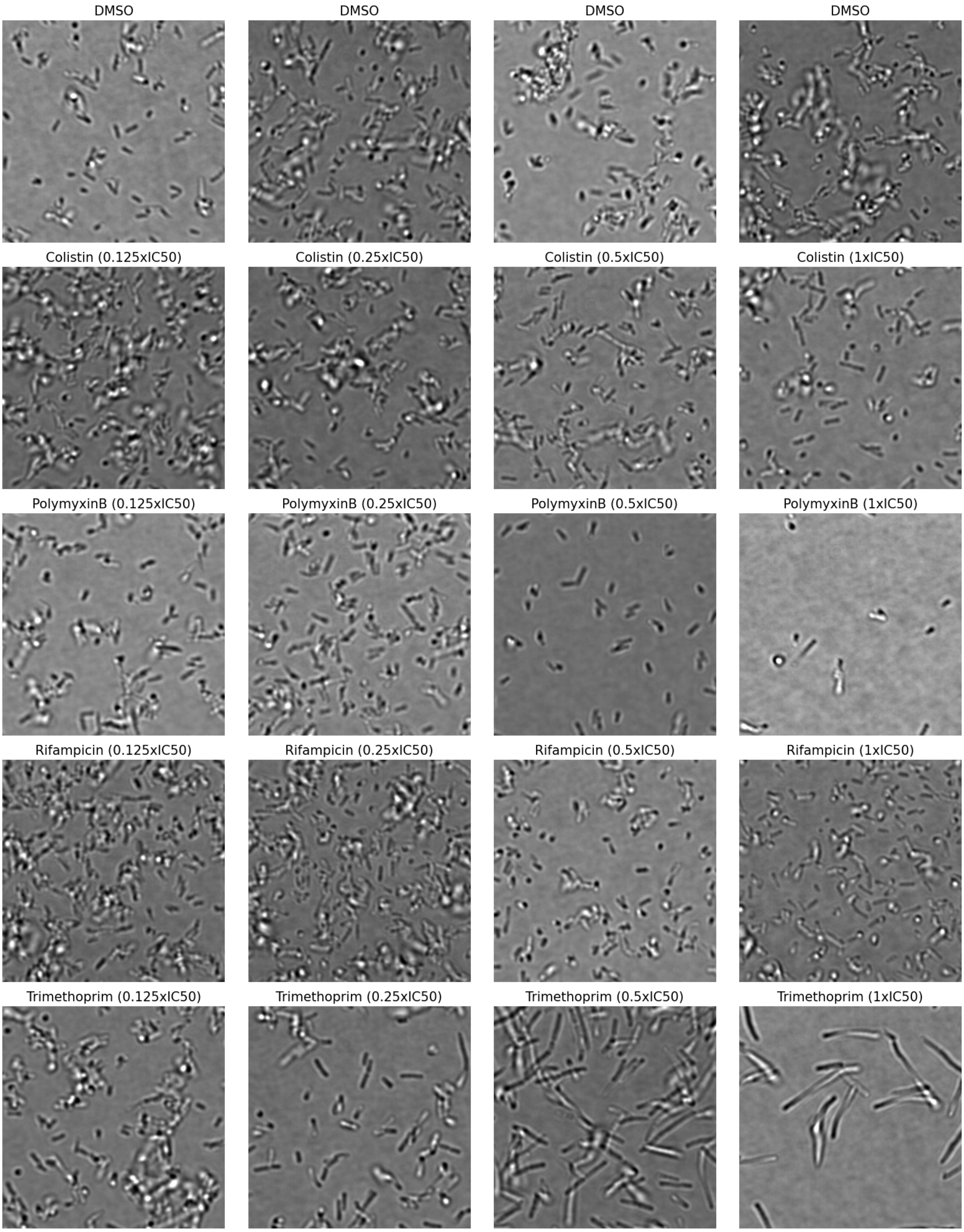

**Supp Fig. 2f: Image gallery for membrane integrity, RNA polymerase and DNA synthesis inhibitors.**

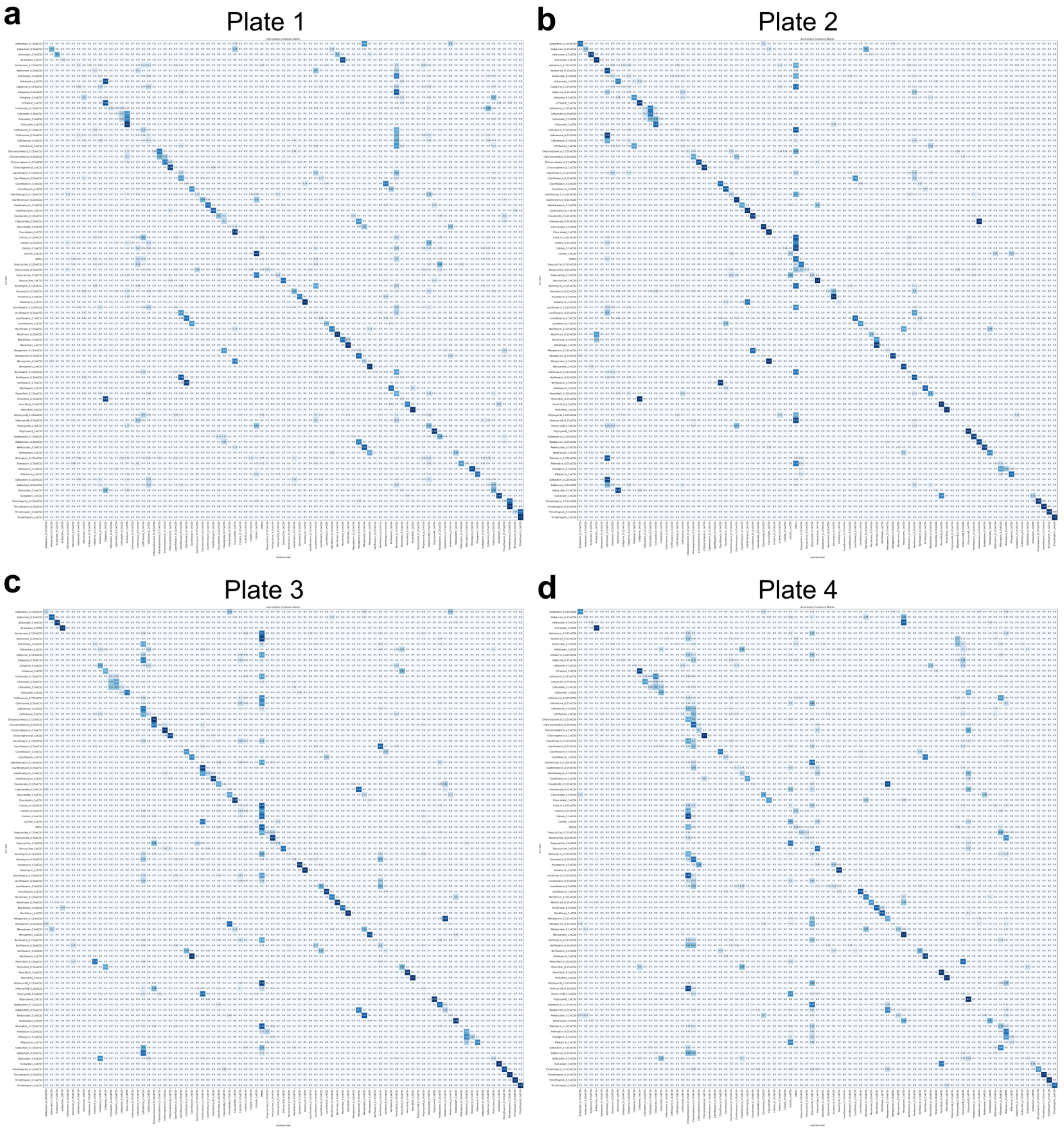

**Supp Fig. 3: Confusion matrices for treatment classification.**

The 89×89 confusion matrices show the predicted treatment combinations (compound identity + concentration = 22 × 4 combinations + 1 untreated control condition) compared to the ground-truth for CNNs trained on three plates and tested on a fourth hold-out test plate. (**a**) Plate 1: 34.8% classification accuracy (31-fold improvement vs. random). (**b**) Plate 2: 47.4% classification accuracy (42-fold improvement vs. random). (**c**) Plate 3: 45.2% classification accuracy (40-fold improvement vs. random). (**d**) Plate 4: 31.3% classification accuracy (29-fold improvement vs. random).

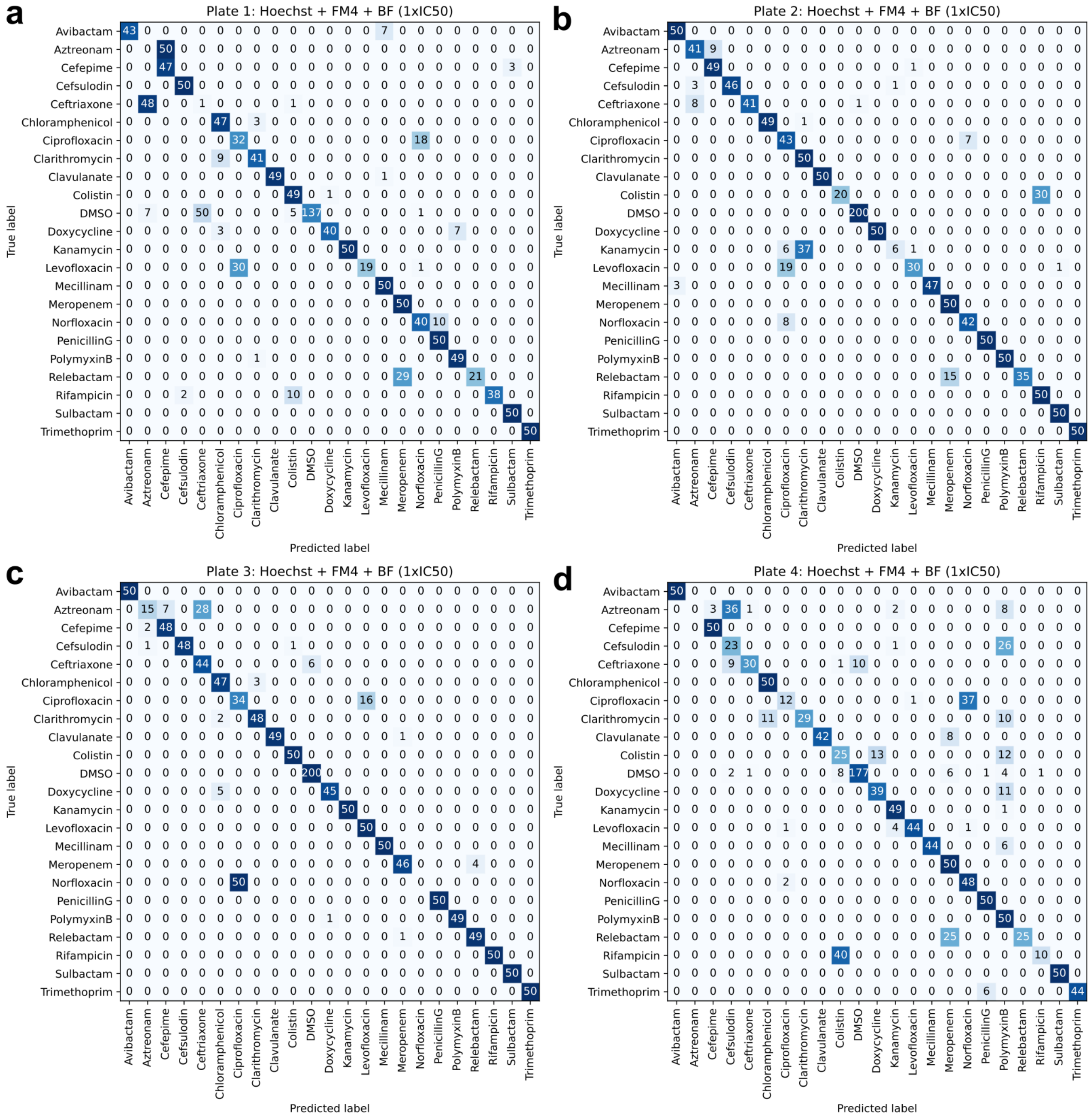

**Supp Fig. 4: Confusion matrices on four hold-out test plates evaluated at $IC_{50}$.**
These 23×23 confusion matrices show the predicted compound identity compared to the ground-truth for CNNs trained on three plates and tested on the fourth. Classification accuracies for the four test plates are as follows: (**a**) Plate 1: 77.15% classification accuracy. (**b**) Plate 2: 88.38% classification accuracy. (**c**) Plate 3: 90.15% classification accuracy. (**d**) Plate 4: 76.23% classification accuracy.

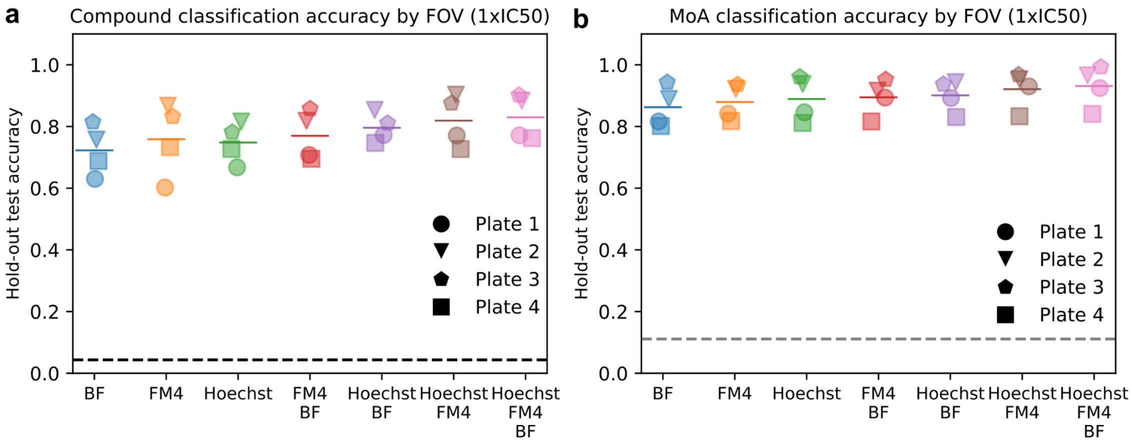

**Supp Fig. 5: Accuracy of compound or MoA classification by FOV for different input channel combinations.**

Models were trained in a cross-validation experiment on data from three plates using different combinations of imaging channels as input and tested on a hold-out test plate with the same combinations of channels at the $IC_{50}$ concentration. Plots show the average per-image classification accuracy for compounds (**a**) and MoAs (**b**). Each symbol corresponds to a distinct plate and horizontal lines show median values. The dashed line indicates random classification.

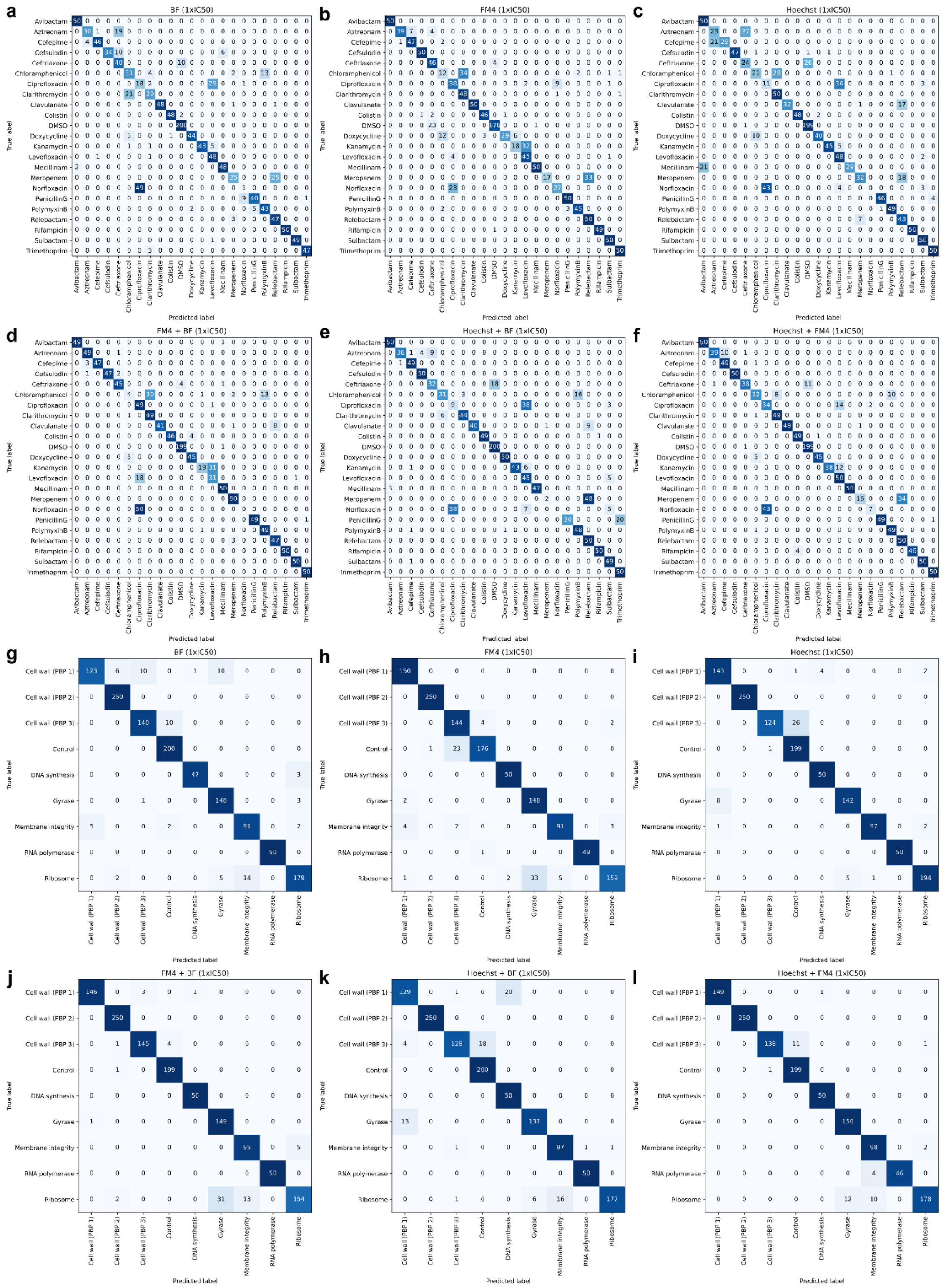

**Supp Fig. 6: Confusion matrices for different channel combinations at $IC_{50}$.** Models were trained in a cross-validation experiment with three plates using different combinations of imaging channels as input and tested on a hold-out test plate at $IC_{50}$ with the same imaging channels. Here, confusion matrices for one hold-out test plate are shown. Confusion matrices by compound (**a-f**) and by MoA (**g-l**) are shown for different input channel combinations as indicated on top of each matrix. Confusion matrices for a model trained and tested on the same hold-out plate for all three colour channels are shown in **Fig. 2b** and **2c**.

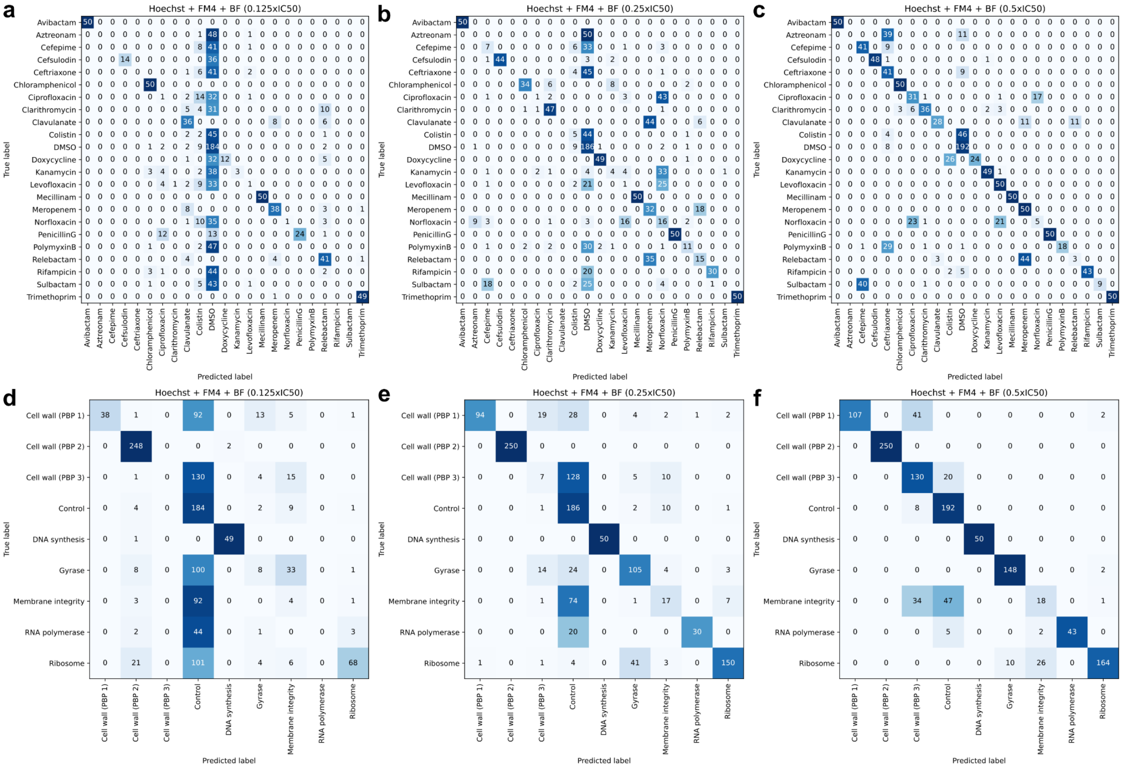

**Supp Fig. 7: Confusion matrices for sub-IC$_{50}$ concentrations.**
Confusion matrices evaluated at different sub-IC$_{50}$ concentrations (0.125×IC$_{50}$ to 0.5×IC$_{50}$) are shown by compound (**a-c**) and by MoA (**d-f**) showing increasing classification accuracy with increasing concentration. All three colour channels were used.

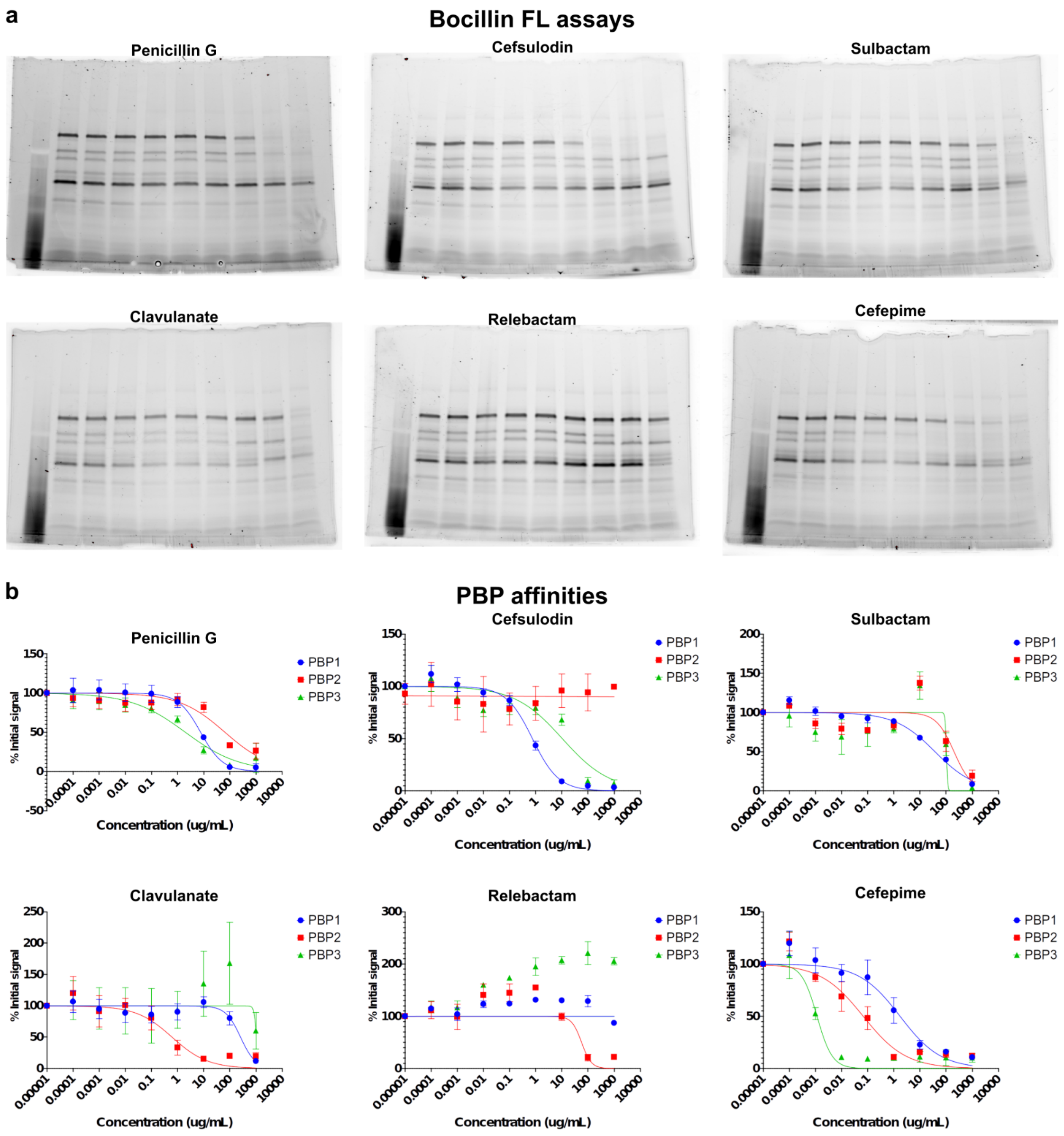

**Supp Fig. 8: Identifying PBP specificities with Bocillin FL competition assays.** (**a**) The fluorescent penicillin analogue Bocillin FL labels PBPs in *E. coli* as resolved by SDS-PAGE. Using a competition assay against a dilution series of PBP inhibitors, changes to the Bocillin FL labelling of each individual PBP can be quantified to reveal the specificity of these inhibitors. (**b**) The quantified PBP bands were plotted against antibiotic concentrations at increasing concentrations.

**Supp. Table 1: PBP specificities from literature.**

| Antibiotic | PBP specificity | PBP1A $IC_{50}$ | PBP1B $IC_{50}$ | PBP2 $IC_{50}$ | PBP3 $IC_{50}$ | Unit | *E. coli* Strain | Ref. |
|---|---|---|---|---|---|---|---|---|
| Cefsulodin | 1 | 0.47 | 3.7 | >250 | >250 | µg/mL | DC0 | (1) |
| Penicillin G | 1 | 0.5 | 3 | 0.8 | 0.9 | µg/mL | DC0 | (1) |
| Sulbactam | 1 | 32 | 1024 | 128 | 512 | µg/mL | J62-1 | (2) |
| Avibactam | 2 | $>10^4$ | 310 ± 460 | <100 | $>10^4$ | µM | MG1655 | (3) |
| Mecillinam | 2 | >250 | >250 | <0.25 | >250 | µg/mL | DC0 | (1) |
| Meropenem | 3 | $>10^4$ | 110 ± 86 | 0.01 ± 0.00 | 0.2 ± 0.2 | µM | MG1655 | (3) |
| Clavulanate | 2 | 128 | 256 | 8 | 256 | µg/mL | J62-1 | (2) |
| Relebactam | - | - | - | - | - | - | - | - |
| Aztreonam | 3 | 970 ± 940 | 1.6 ± 0.9 | $>10^4$ | 0.09 ± 0.09 | µM | MG1655 | (3) |
| Cefepime | 3 | 5.5 ± 2.6 | 2.6 ± 2.4 | 0.2 ± 0.1 | $<10^{-4}$ | µM | MG1655 | (3) |
| Ceftriaxone | 3 | 4.9 ± 2.2 | 0.2 ± 0.2 | 1.5 ± 0.9 | 0.001 ± 0.003 | µM | MG1655 | (3) |

**Supp. Table 2: Antibiotic source**

| Antibiotic | Product name | Product supplier | Catalog number |
|---|---|---|---|
| Cefsulodin | Cefsulodin Sodium Salt | MP Biochemicals | 098677 |
| Penicillin G | Penicillin G sodium salt | Sigma | 13752 |
| Sulbactam | Sulbactam | Sigma | PHR2576 |
| Avibactam | Avibactam sodium | RayBiotech | 332-11720-1 |
| Mecillinam | Mecillinam | Cayman Chemicals | 9002008 |
| Meropenem | Meropenem trihydrate | Sigma | PHR1772 |
| Clavulanate | Lithium clavulanate | Sigma | L0720000 |
| Relebactam | Relebactam | MedChemExpress | HY-16752 |
| Aztreonam | Aztreonam | MP Biomedicals | 150415 |
| Cefepime | Cefepime hydrochloride | Sigma | PHR1763 |
| Ceftriaxone | Ceftriaxone disodium salt hemi(heptahydrate) | Sigma | C5793 |
| Chloramphenicol | Chloramphenicol | Sigma | C0378 |
| Clarithromycin | Clarithromycin | Sigma | PHR1038 |
| Doxycycline | Doxycycline hyclate | Sigma | D9891 |
| Kanamycin | Kanamycin sulfate from Streptomyces kanamyceticus | Sigma | K4000 |
| Ciprofloxacin | Ciprofloxacin hydrochloride | Euromedex | UC5074-B |
| Levofloxacin | Levofloxacin Anhydrous | Sigma | 28266 |
| Norfloxacin | Norfloxacin | Sigma | N9890 |
| Rifampicin | Rifampicin | Sigma | R3501 |
| Trimethoprim | Trimethoprim | Sigma | T7883 |
| Colistin | Colistin Sulfate | Sigma | PHR1605 |
| Polymyxin B | Polymyxin B sulfate salt | Sigma | P4932 |
